# The representation of estimated mood in primate anterior insular cortex

**DOI:** 10.1101/2025.04.22.650010

**Authors:** Nicole C. Rust, You-Ping Yang, Catrina M. Hacker, Veit Stuphorn

## Abstract

Understanding how the brain reflects and shapes mood requires resolving the disconnect between behavioral measures of mood that can only be made in humans (typically based on subjective reports of happiness) and detailed measures of brain activity only available in animals. To achieve this, we developed a mood model to predict behavioral fluctuations in human subjective happiness as individuals experienced wins and losses during a gambling task. Next, we investigated how this operationalization of mood was reflected behaviorally and in the brains of two monkeys engaged in the same gambling task. We found a remarkable alignment between human mood model signatures, the impact of estimated mood on monkey choice, and the persistent responses of units in monkey anterior insular cortex — including a matched timescale of integration across events. In comparison, the same signatures were only weakly reflected in lateral prefrontal cortex, suggesting that insular mood representations do not trivially follow from a signal broadcast to all higher brain areas. These results are consistent with a model in which the brain transforms experiences into mood by integrating events through a recurrently connected network of excitatory and inhibitory pools of neurons. These are among the first detailed insights into the nature of putative mood representations in the primate brain.

## Introduction

Of all the functions of the brain and mind, mood is among the most mysterious. In comparison with acute emotions that are often targeted at something specific (like the fear of a tiger), moods are typically more diffuse and slowly changing, as mood reflects an integration across multiple events (1–5). Understanding mood is important, given the vast unmet need of individuals with mood disorders including major depression and bipolar disorder. A remarkable 21% of adults will experience a mood disorder at some point in their lives (6), and a third of those will be resistant to existing treatments (7). The development of new and improved behavior-based and brain-based interventions for mood disorders could benefit from a better understanding of how mood is supported and shaped by the brain (8, 9).

To date, researchers have learned a considerable amount about how mood is influenced by what happens at the highly detailed molecular level (e.g. by serotonin neurotransmission, (10)). They’ve also made progress toward understanding the broader scale brain networks involved in mood via fMRI (e.g., 10). In comparison, remarkably few studies have focused on how patterns of brain activity exhibited by populations of individual neurons drive the experience of mood, how processing in the brain converts external and internal experiences into those brain activity patterns, and how these mood representations influence behavior. To fill this gap, we will need to overcome impediments that make mood more challenging to study than other brain functions such as perception, memory, and decision making.

Many of the challenges associated with understanding mood follow from the fact that mood is a subjective experience for which there is no external ground truth — no one knows how happy an individual is but themselves. Consequently, mood cannot be studied by benchmarking against performance on objectively defined tasks in the same way many other brain functions are studied. Likewise, mood is difficult to study in animals because we lack good ways to measure how they feel (2). While behavioral tests such as the forced swim test have been used to study mood and depression in animals, their relevance to understanding mood in humans is controversial (12). Similarly, inferring mood from innate behaviors such as facial expressions is thought to be problematic (13, 14).

At the same time, the types of densely sampled, high spatial and temporal resolution measures of neural activity that are best suited to investigating neural representations are difficult to obtain in humans. While meaningful data can be gathered from human patients with electrodes implanted for epilepsy or other clinical purposes, those measurements are often sparse and are typically lower-resolution measures of activity that may not reflect the action potentials that the brain uses to communicate within and between brain areas (15). Similarly, while a handful of studies have made notable progress in “decoding mood” from human neural activity (16–18), all have taken a clinically inspired engineering approach focused on the accuracy with which mood can be decoded from the human brain as opposed to focusing on the nature of the neural code for mood itself.

Here, we sought to overcome these challenges by applying an approach that seeks to understand mood by using models to link across data of different types. The links between the type of happiness fluctuations we study here and those that go awry in mood disorders like depression have yet to be established (19). However, given the paucity of understanding of how the different dimensions of mood are reflected in the brain, descriptions of neural mechanisms that shape typical happiness fluctuations may be an important foundational step toward the longer-term goal of understanding mood disorders. Such is the basic premise of fundamental neuroscience research (20) that has led to transformative breakthroughs in studying brain functions like vision, memory, decision making and motor control.

## Results

To study mood and its neural correlates in a rigorous way, we need to operationalize its definition, measure it, and systematically modulate it. Our operationalization and modulation of mood captures four generally well-agreed-upon characteristics of it: 1) measures of mood often probe subjective reports of affect (21–23); 2) mood reflects an integration of events as opposed to individual experiences (1, 5, 24–29); 3) mood is modulated bidirectionally by both positive and negative events (1, 4, 5, 21, 22, 26, 29); and 4) mood influences cognitive processing such as the propensity to engage in risk (21, 29–34). Below, we refer to the operationalization we use here as “mood” for ease of reading (as opposed to the implication that mood is a singular construct, driven by a single process).

Our approach builds on the work of Rutledge et al. (35), who introduced a way to model and thus predict how an individual’s mood will fluctuate in the context of a gambling task. As an overview, they modulated mood via monetary wins and losses on gambling trials, and they measured mood with subjective happiness probes inserted every few trials. They then used this behavioral data to fit a model that predicted mood based on the wins and losses each individual experienced, and that model provided reasonable accounts of subjective happiness fluctuations.

Characteristic of mood, Rutledge et al. (35) found that individuals’ answers to the question, “How happy are you right now?” depended not just on events on the last trial but an integration of events across ∼7 trials, on average across subjects. They also determined via fMRI that this operationalization of mood was reflected in anterior insular cortex (AIC). In comparison, fMRI activation of the striatum was correlated with gambling trial outcomes but not while answering questions about happiness itself. These results are consistent with multiple lines of evidence that tie AIC to the conscious experience of affective feelings, including mood (36). However, they do not answer the question of how happiness is reflected in this brain region at the level of individual AIC neurons, nor do they shed insight into how mood is computed by the brain.

For answers, we analyzed one of the few existing datasets of AIC activity in the nonhuman primate brain, measured at single-unit resolution. In this study (37), rhesus monkeys performed a gambling task similar to that investigated by Rutledge et al. (35). As an overview of our report, we begin by developing a class of mood models to predict human behavioral mood modulation. Next, we confirm that this subject-generalized mood model is relevant to monkey behavior and then describe its neural correlates in the monkey brain, focusing on AIC following the fMRI evidence cited above. Here, we assume that the representation of human subjective happiness can be meaningfully studied by probing neural representations in monkeys, contingent on finding straightforward and robust behavioral and neural correlates.

We found a remarkable alignment between the mood behavioral signatures derived from human subjective accounts, the impact of estimated mood on monkey choice, and the responses of individual AIC units during the behaviorally unconstrained period between trials. These mood representations were reflected in the activity of a large fraction of the AIC population. In comparison, the same mood signatures were nearly absent in lateral prefrontal cortex. This nontrivial alignment between human behavioral estimates of happiness, the impact of estimated mood on monkey choice and monkey neural activity could provide a putative neural correlate of mood that can be built upon to meaningfully understand how the brain generates mood.

### Human behavioral mood models

Our method for determining the neural correlates of mood fluctuations combined the approaches developed by Rutledge et al. (35) with the details of the gambling task performed by the monkeys during AIC recording, originally reported by (37). In the latter study, the monkeys gambled for wins and losses of 0-3 tokens per trial, and they received a liquid reward when the number of tokens reached 6 (Fig 1A). Specifically, on each trial the monkeys selected between a certain outcome and a gamble between two options weighted by different probabilities, resulting in a change in token number that ranged from −3 (lost) to 3 (gained) on each trial. On some trials, the best choice was obvious, such as a +3 token certain reward versus a +3/0 token gamble with 0.1/0.9 probabilities (where selecting the certain reward was the obvious best choice). On other trials, the best choice was less obvious, such as −1 tokens for certain versus a 0/-3 gamble with 0.5/0.5 probabilities. When the total number of tokens reached 6, the monkey received a juice reward and 6 tokens were subtracted from the token total.

**Figure 1.**
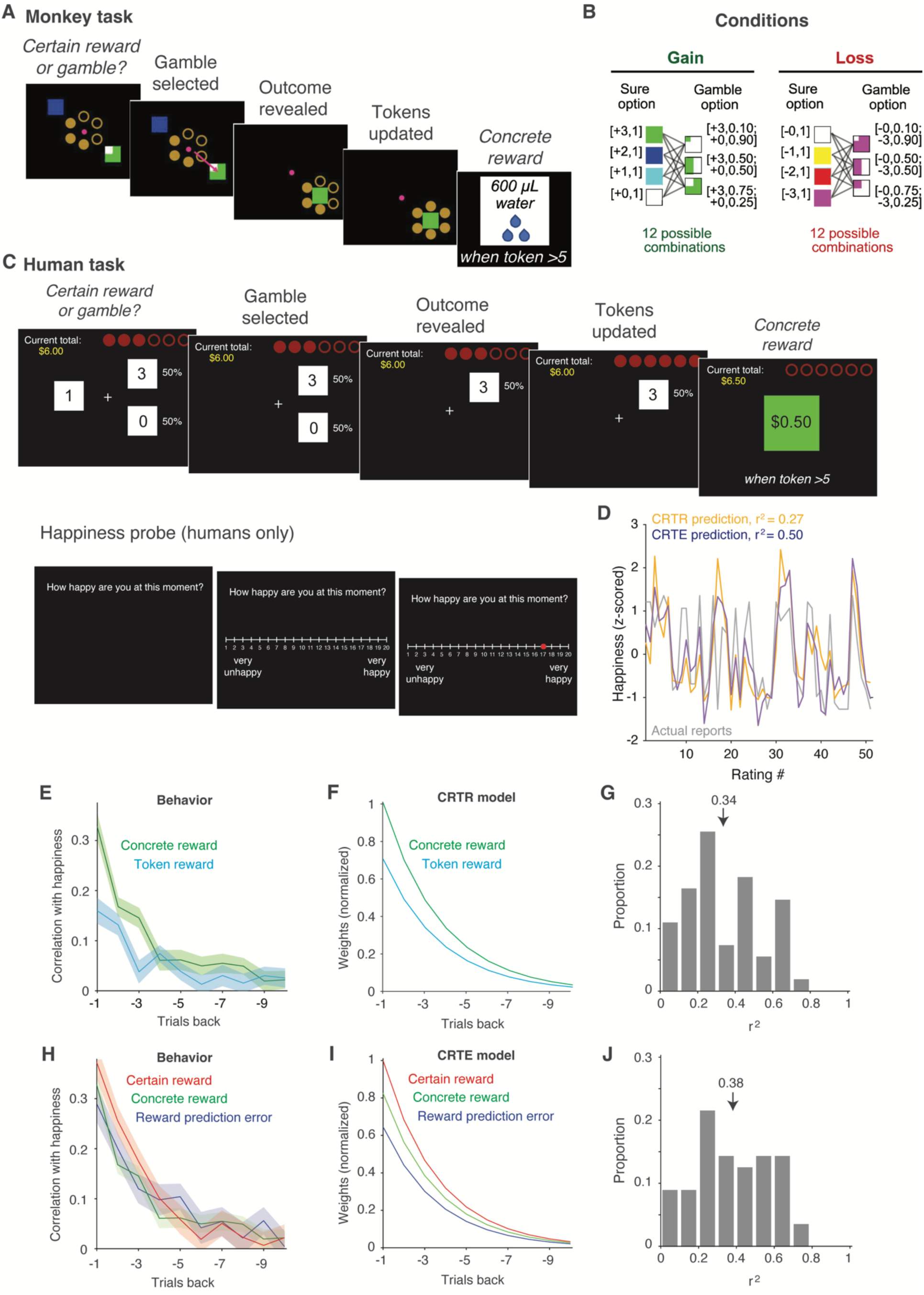
*Behavioral paradigm and mood model predictions.* A) On each trial of the monkeys’ gambling task, monkeys selected between a certain reward (upper left) and a gamble (lower right) for a token reward where tokens were reflected as colored blocks that the monkeys had learned to map to their token values. On each trial, −3 to 3 tokens were rewarded. When the token number reached 6, the monkeys received a liquid reward. B) Experimental conditions and their symbols. Trial types were parsed into token wins (Gain) and token losses (Loss) trials, with all possible combinations of 4 certain rewards and 3 gambles for each type. C) The human version of the task was designed to match the monkeys’, with the exception that token numbers rather than symbols reflected the options, and humans won money as opposed to water. A happiness probe was interleaved every 2-3 trials. D) The happiness reports of a representative human subject (gray), along with the subject-generalized happiness model prediction of how their happiness would fluctuate as computed by the CRTR model (yellow) and the CRTE model (purple). E) Correlation between human happiness reports and both concrete and token rewards, demonstrating the influence of both on happiness reaching back ∼6-8 trials. Lines indicate means and the shaded region indicates standard error across 55 subjects. F) To compute CRTR model predictions for the subject shown in panel D, the monetary and token reward events that happened during the course of the experiment were integrated with the weights shown, determined by fitting the model to the other 54 subjects’ data. Compare with panel E. G) Distributions of r^2^ across all 55 subjects when each subject was held out and the CRTR model was fit to the remaining 54 subjects’ data. Arrow indicates the mean (0.34). H) Correlation between human happiness reports with: concrete reward, certain reward (tokens won on gambling trials when the certain option was selected) and reward prediction error when the gambling option was selected (with the same conventions as Fig 1E). Reward prediction error was computed as the difference between tokens won and expected value, where expected value was defined as the average token wins on gambling trials up to the trial in question. Because expected value has minimal trial-by-trial modulation to correlate with, it (the final component of the CRTE model) is not shown. I) Integration weights for the CRTE model with the same conventions as panel F. J) Distribution of r^2^ across all 55 subjects for the CRTE model. Arrow indicates the mean (0.38).

Trials came in two types, parsed by token wins (gains) and losses; altogether, there were 12 types of gain trials and 12 types of loss trials (Fig 1B). In addition, each of the 13 options (both certain rewards and gambles) was presented independently on “forced choice” trials in which the monkey was required to select that option. In total, 37 unique trials were randomly shuffled and each presented once before re-randomization.

The human version of the task was, by design, intended to match the monkeys’, with the exception that humans received a $0.50 reward each time the number of tokens reached 6 instead of liquid (Fig 1C). For ease, we refer to both liquid (monkeys) and money (humans) as “concrete” rewards to differentiate them from the more abstract “token” rewards that were accumulated to receive concrete ones. In addition, a mood probe was introduced in the human version of the task every 2-3 trials where they were asked to rate their momentary happiness on a scale of 1-20 (Fig 1C; Fig 1D gray). The mean and std of the happiness range used by subjects was 11.8 +/− 5. (Supplementary Fig 1).

Here we begin by demonstrating the relationship between human subjective happiness reports and the two types of rewards (concrete and token), not just received on the last trial but across a handful of trials. Consistent with the notion that mood reflects an integration of events (as well as the previous results by Rutledge et al. (35)), reports of subjective happiness were positively correlated with concrete rewards across ∼8 trials leading up to a happiness probe on average, across the 55 human subjects (Fig 1E, green). Happiness was similarly correlated with token reward across the last ∼6 trials (Fig 1E, cyan).

Next, we evaluated several different mood models for their ability to predict happiness fluctuations. To focus on what modulates reward in the gambling task (as opposed to the external factors that set its baseline), happiness reports were z-scored for each subject. Consistent with the intuitions depicted in Fig 1E, one conceptually straightforward model computed subjective reports of happiness as a weighted sum of concrete and token rewards integrated with a fading average over the last ∼7-8 trials. We will refer to this as the “concrete reward and token reward (CRTR) model”. To compute CRTR happiness model predictions for the subject shown in Fig 1D, the events this individual experienced during the gambling task were combined as a weighted sum of terms capturing concrete and token rewards, with the relative weightings over time shown in Fig 1F (see Supplementary Fig 2 for concrete and token reward traces). To determine these weights, the model was fit to data from 54 human subjects and then used to predict this individual’s held-out data. Shown in Fig 1D are the happiness fluctuations for one representative human subject (gray), along with the predictions of this model (yellow). For this subject, the CRTR subject-generalized model predicted many (but not all) of the features of how this person’s happiness would fluctuate across the gambling task based on the wins and losses they experienced (Pearson’s r^2^ = 0.27).

Building on this intuition of how the CRTR happiness model operates, we extend it to the other models we considered. Across the population, models with fewer terms (such as concrete or token rewards alone) produced worse happiness predictions than the CRTR model (Supplementary Table 2). Conversely, a subset of models that parsed token-related events into additional terms provided better predictions (Supplementary Table 2). Among all the models, four produced similar, highest subject-averaged predictions. What set these models apart from the others was that they differentially weighted concrete rewards versus token rewards and they deconstructed token rewards into a weighted sum of at least two terms — one for tokens won on trials when the certain reward option was selected, and one (or more) terms for tokens won on trials when the gambling option was chosen (Supplementary Table 1). Given the similarity of these four model predictions, coupled with the similarities in the parameterization of these models, we hesitate to declare that there is a singular “best model” that accounts for subjective happiness reports in this data. Rather, a more parsimonious description is that a class of models provides a decent description of the data overall, and these are ones in which happiness depends on a weighted combination of concrete rewards and token-related events parsed into certain reward and gambling option selections.

To illustrate the properties of this class of models, we focus on one that we refer to as the Concrete Reward Token Expectation (CRTE) model. The CRTE model estimated happiness based on concrete rewards (Fig 1I, green) and token-related events parsed into 3 terms. Those included one term that captured tokens received when the certain reward option was selected (Fig 1I, red). In addition, two terms captured tokens received on gambling trials, including one to capture expected value (defined as the average token wins on gambling trials up to the trial in question, thus leaving minimal trial-by-trial modulation to correlate with; not shown). The second captured the difference between the expected value and the token amount won (reward prediction error, Fig 1I, blue). Like the terms for the CRTR model, the components of the CRTE model correlated with happiness reports across ∼6 trials (Fig 1H), and the model weights fit for those terms bore a striking resemblance to correlations in the raw data (Fig 3I). For the subject presented in Fig 1D, model predictions were higher for the CRTE (Fig 1D purple, Pearson’s r^2^ = 0.50) as compared to the CRTR model (Fig 1D yellow, Pearson’s r^2^ = 0.27). Across the 55 subjects, average CRTE r^2^ was significantly higher than average CRTR r^2^ (mean CRTR: Pearson’s r^2^=0.34, Fig 1G; mean CRTE Pearson’s r^2^=0.38, Fig 1J; one-way paired t-test, t(54)=3.33, p=0.002).

As the distributions of r^2^ for both models reflect, the predictions of these models were better for some individuals than others. Focusing on the CRTE model, this account produced exceptional predictions for some individuals (for examples, see Supplementary Fig 3, top). In comparison, the subjective happiness reports of a handful of individuals were insensitive to the events of the gambling mood induction paradigm, and consequently, models based on those events provided poor predictions of happiness fluctuations (for examples, see Supplementary Fig 3, bottom). Eliminating subjects like these did not qualitatively change the results we report here (data not shown). In all the analyses presented in this report, the data from all 55 subjects are included.

### Estimated happiness increases risk seeking and decreases loss aversion in monkeys

To recap, the CRTR model fit to subjective happiness reports during this gambling mood induction paradigm computes mood based on fluctuations in concrete and token rewards, computed as a fading integration across the last ∼7 trials (Fig 1F). Models that deconstruct token rewards into additional terms (like the CRTE model) perform moderately but significantly better (Supplementary Table 2). To understand the neural correlates of mood, we looked to determine how these behavioral signatures were reflected in high-resolution measures of AIC neural activity recorded in the rhesus monkey brain.

We hypothesized that the monkey brain computes and reflects the same signatures that contribute to human subjective happiness reports, and one locus of that representation is AIC. Notably, our hypothesis is restricted to the nature of neural representations in the monkey brain (which we can measure) while remaining agnostic about whether the monkeys subjectively experience happiness (which we have no way to access). However, analogous to other measures of affect in humans (38) and animals (25), we could assess whether the estimated mood modulations induced by the gambling paradigm had a predictable impact on the monkeys’ behavior, including their risk seeking and the urgency of their responses (reflected in their reaction times).

To perform these analyses, we estimated monkey happiness by passing the monkeys’ trial options and their outcomes through the subject-generalized mood models fit to humans (number of trials M1 = 77469; M2 = 25544). We then determined whether the monkeys’ choice to gamble was influenced not only by current trial details (including the expected values of the two options before them and the number of tokens at trial start), but also by happiness estimated by the CRTE model, computed based on the events leading up to a trial. For both monkeys, the probability of selecting the risky (gamble) option increased as estimated happiness increased on both gain and loss trials (regression analysis, based on a coding of gambling choice as 0/1; gain trials: M1 β = 0.004, p = 2*10^−7^; M2: β = 0.003, p=0.014; loss trials: M1 β = 0.002, p = 0.001; M2: β = 0.005, p=0.0001). From the perspective of frameworks that separately consider choice behavior for wins versus losses, these results are consistent with happiness increasing “risk seeking” and reducing “loss aversion”.

Likewise, reaction times (relative to the onset of the target option) significantly changed with happiness on gain trials (regression analysis, mean/sd RT across M1 and M2 = 247+/−80 ms; M1 β = 2.16, p = 1*10^−65^; M2: β = 1.53, p= 3*10^−14^). On loss trials, reaction times significantly decreased with increasing estimated happiness in one monkey, with the same trend but not a significant relationship in the other M1 β = −0.27, p = 0.10; M2: β = −1.40, p=1*10^−12^).

To better understand how happiness influenced the monkeys’ choice behavior, we extended a model based on the prospect theory of decision making (39) previously used to describe choice behavior in these monkeys (37). Prospect theory captures notable signatures of risk-taking, including that individuals 1) are more sensitive to changes in value for losses versus wins (here: tokens on gain versus loss trials, Fig 1B), and that they 2) make decisions about risk-taking based on their current wealth (here: accumulated token count, Fig 1A). In the previous report, both signatures were reflected in the gambling behavior of these monkeys (37). In the case of wealth, the probability that the monkeys would gamble depended on their current token assets but differed for trial type: they were more likely to gamble as assets increased on gain trials and less likely to gamble as assets increased on loss trials. Differences between gain and loss trials were captured in the behavioral model as separate terms for risk seeking and loss aversion.

Beginning with this choice model (of when the monkeys would gamble based on the options before them and current token assets), we looked to see whether extending the model to incorporate estimated happiness (computed based on the events leading up to a trial) would provide an even better account of those choices. Models were evaluated via cross-validation by comparing performance on held-out data (described here, see Methods) as well as models fit to all data (Supplementary Table 3). For both monkeys, adding two parameters to account for modulation by happiness as estimated with the CRTE model (one for risk seeing and another for loss version) significantly increased model predictions over a model without happiness included (one-way paired t-test M1: t(4)=7.70, p=0.002; M2: t(4)=12.0, p=0.0003). In the model, higher happiness led to more risk-seeking behavior and less loss-averse behavior in both monkeys (Fig 2; Supplementary Table 3). For the example (gain) trial shown, modulations of the probability of selecting the gamble from the theoretically lowest to highest possible happiness were large, increasing 41% to 69% in monkey 1, and from 53% to 91% in monkey 2.

**Figure 2.**
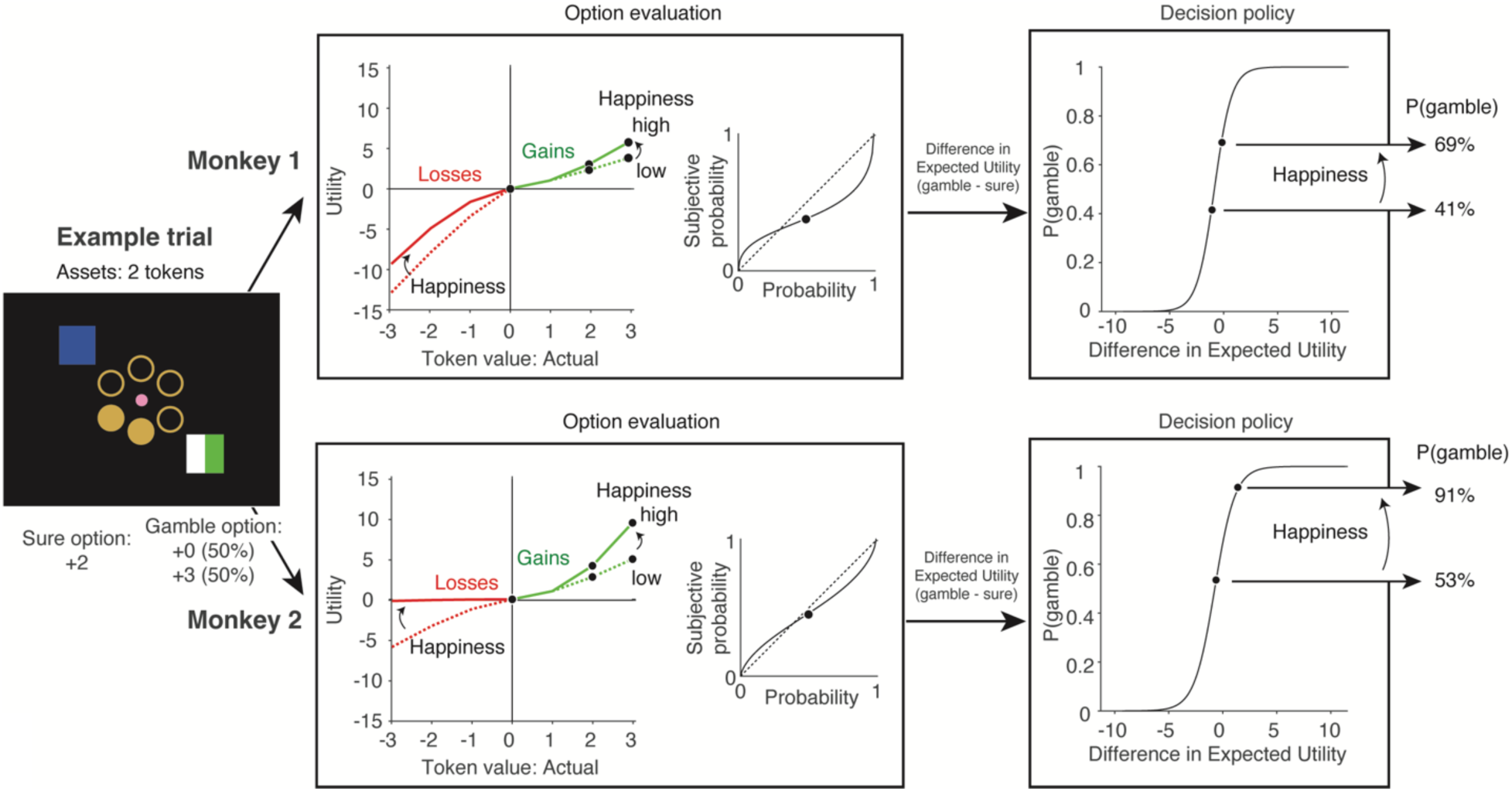
*A model of how happiness influences the monkeys’ decisions to gamble.* Here, a prospect theory model of decision making was extended to incorporate happiness as estimated by the CRTE model. Shown are the models fit to data from monkey 1 (top) and monkey 2 (bottom), along with 1 example trial. On this trial, the monkeys began with 2 token assets, and they had to select between winning 2 tokens for certain versus a gamble between winning 0 or 3 tokens with 50/50% probability (the model input; left). The model output reflects the probability that each monkey will choose the gamble option (right), computed for lowest and highest possible estimated happiness (1 and 20, respectively). In the “Option evaluation” component of the model, token value is converted into the subjective quantity utility (left), determined separately for gains (green) and losses (red) by a combination of baseline terms and modulations of those terms by token asset (see Methods). In addition, the conversion is influenced by happiness (dashed lines: low happiness; solid lines: high happiness; see Methods). Shown is the conversion for the lowest and highest possible happiness values for trials that begin with 3 token assets. Black dots correspond to the token values under consideration in the example trial (0, 2 and 3). Next, utility is combined with a function that converts the probability of outcomes into a subjective probability to compute expected utility (middle, black solid line; dashed line indicates identity for reference). In the “Decision policy” component of the model (left), the difference in expected utility for the sure option versus the gamble is passed through a sigmoidal function to determine the probability of selecting a gamble. As shown, a change in happiness from 1 to 20 modulates the model-predicted probability of selecting a gamble on this example trial from 41% to 69% in monkey 1 and from 53% to 91% in monkey 2.

Similar results were obtained when happiness was estimated with the CRTR model, including that a choice model with two happiness terms performed better than one with no terms (one-way paired t-test M1: t(4)=5.84, p=0.004; M2: t(4)=11.31, p=0.0004). In addition, the choice model with happiness estimated by the CRTE model performed better than the same model with happiness estimated by the CRTR model (one-way paired t-test M1: t(4)=4.98, p=0.008; M2: t(4)=8.66, p=0.001).

Together, these results establish that estimated happiness is a quantity that affects the monkeys’ behavior, thereby supporting the relevance of looking into how it is computed by the monkey brain.

### Insight into the neural mechanisms that shape mood

Following on the fMRI results of Rutledge et al. (35) that answers to the question, “How happy are you right now?” are reflected in anterior insular cortex (AIC), we turned to look for single neuron correlates of mood there. Building on the behavioral observation that subjective happiness reflects an integration of recent events, we suspected that mood might be reflected in AIC via the type of persistent, elevated activity reported in other brain regions where representations are often distributed across individual, persistent units (40). Because mood is an affective state that changes over a timescale much longer than a single trial, we reasoned that it should be reflected not just during gambling trials, but also between them. We thus probed the time period *in between* trials of the gambling task for mood representations. A component of our hypothesis is the proposal that mood is reflected in AIC in the absence of querying it. In other words, we hypothesize that the answer to the question, “How happy are you right now?” is reflected in AIC even when the question is not asked. Should this not be true, we would fail to find a strong neural correlate there.

To test this hypothesis, we focused on the CRTE model, which was among the best performing in terms of its ability to account for human subjective reports. It also illuminates the central signatures of mood captured by the class of best models: that happiness follows from 1) an integration across different types of rewards (concrete rewards and token rewards, with the latter parsed into certain reward selections versus gamble selections leading to reward prediction errors) and 2) an integration across trials. Any putative neural correlate of mood would need to reflect these same signatures. To test the hypothesis that these are reflected in AIC, we computed estimated happiness by passing the monkeys’ trial options and their outcomes through the subject-generalized CRTE mood model fit to humans. The responses of one remarkable AIC neuron are shown in Fig 3A, illustrating how the activity of this neuron persists throughout the 1s period before trial onset in a manner that reflects estimated happiness. Fig 3B shows a strong and highly significant correlation between estimated happiness and the firing rate of this individual unit between trials (Pearson’s r(1307)=0.53; p=3*10^−96^), where every dot reflects the spike count in a single intertrial interval period. Fig 3C shows the correlation between this unit’s firing and the components of reward (concrete, certain token reward and token reward prediction error) as a function of trial history, illustrating an integration over ∼7 trials (compare with 1H). In sum, during the period between trials of the gambling task, this unit reflected all the signatures of human subjective happiness in its persistent responses, including modulation by concrete and both components of token rewards, integrated across several trials. We found many units of this type (elaborated below).

**Figure 3.**
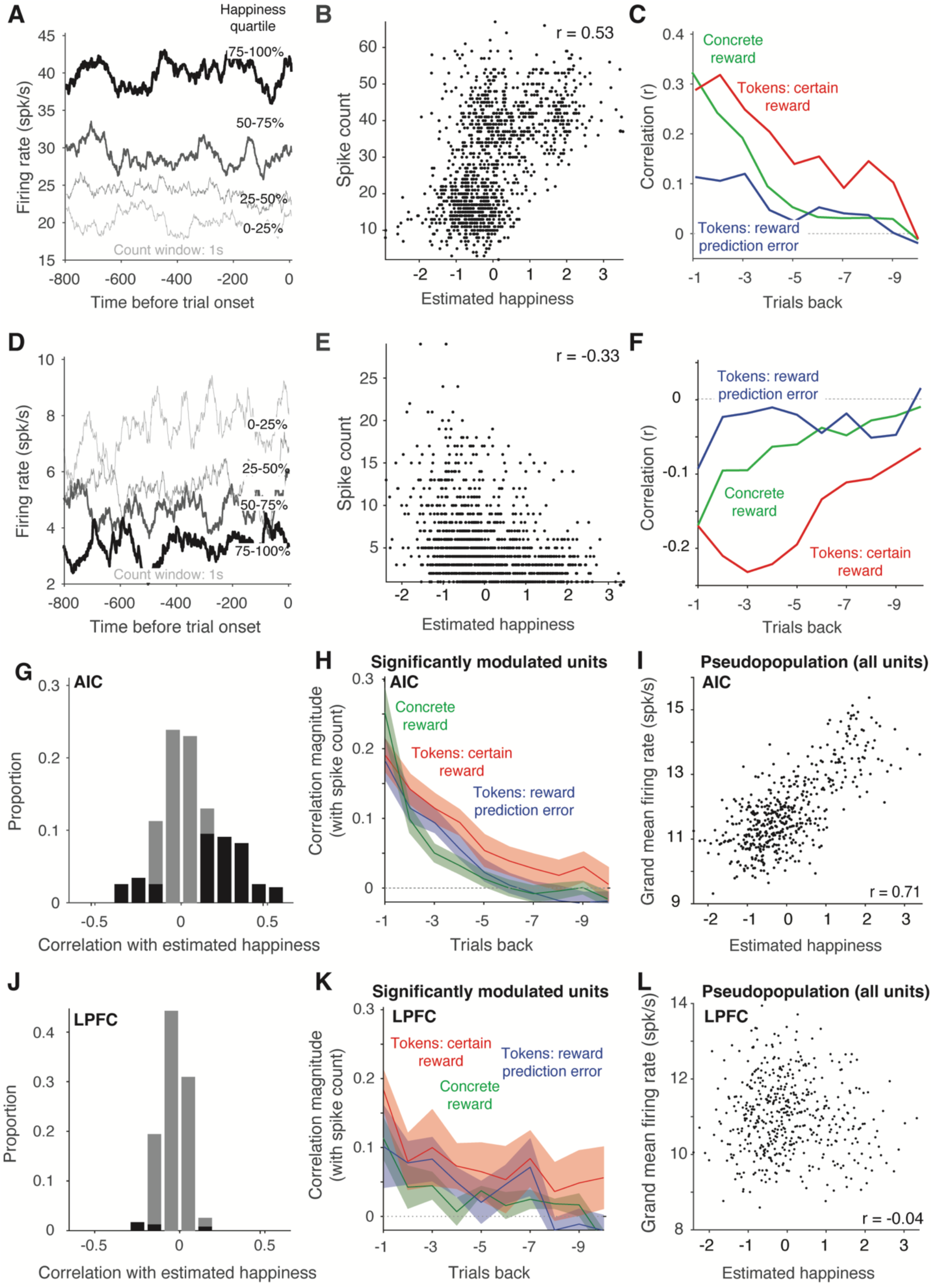
*Individual AIC units reflect happiness estimated by the CRTE model.* A) A PSTH for one monkey AIC unit during the intertrial interval period in the 1s before trial onset, with trials grouped into estimated happiness quartiles. B) For the same unit, the correlation between estimated happiness based on the monkey’s wins and losses and the spike count response, where each dot reflects 1 intertrial interval period. This unit was positively correlated with estimated happiness. C) For the same unit, correlation between the spike count responses, concrete reward history, and token reward history deconstructed into trials in which the certain reward versus the gambling option was selected (thus creating a reward prediction error), computed as described for Fig 1I. D-F) The responses of a second AIC neuron with responses negatively correlated with happiness, with the same conventions as panels A-C. G) Histogram of the Pearson correlation values (e.g., 3B, 3E) across all 229 neurons (gray) and those with significant correlations in black (significance criterion p<0.001). H) Correlation magnitudes (after taking the absolute value of the sign correlation) of spike counts with concrete rewards, certain rewards and reward prediction errors, averaged across all significant units. I) A pseudopopulation was created by aligning units by estimated happiness. Shown is the correlation between estimated happiness and AIC population response vigor. J-L) The same analyses described in panels G-I applied to 224 units recorded in LPFC. For the analyses in G-L, units were limited to 480 trials via equal spacing across the sampled estimated happiness range to equate between them.

While less frequent, we also found AIC units whose responses were strongly negatively correlated with estimated happiness. One example is shown in Fig 3D-F. Like the unit depicted in Fig 3A, this unit’s firing rates persisted across the 1s period before trial onset (Fig 3D); however, the firing rate of this unit was lower when estimated happiness was higher (Pearson’s r(1276)=-0.33; p=4*10^−34^; Fig 3D-E). Likewise, this unit’s firing rates decreased based on concrete rewards and total token number across the last ∼9 trials (Fig 3F). Additional example cells with a range of estimated happiness correlations are presented in Supplementary Fig 4.

To compare units, we equated their numbers of trials to 480 (see Methods). Across the AIC population, the intertrial interval responses of a notable 41% of insula units were significantly correlated with the CRTE model estimate of happiness (significance criterion: p<0.001). When parsed by positive versus negative correlations, 3.7-fold more units were positive (proportions 32% positive versus 9% negative of the total population, respectively; Fig 3G). Correlation magnitudes averaged across significantly modulated neurons (with the correlation signs flipped for negatively correlated happiness units) reflected an integration across ∼6 trials back (Fig 3H), and bore a striking resemblance with behavior (compare with Fig 1H).

We also assessed how fluctuations of the combined AIC population aligned with the behavioral subject-generalized mood model. In these experiments, the neurons were recorded individually, most often in different sessions. We thus created a pseudopopulation by aligning trials by their estimated happiness similar to the approaches described by (41). The resulting pseudopopulation included all units (not just significant units) and thus contained the responses of 229 AIC units on 480 trials. Correlations between estimated happiness and the overall vigor of the AIC population response were strong and highly significant (Pearson’s r(480)=0.71, p=8*10^−75^; Fig 3I), and this was true in each monkey individually (monkey 1: Pearson’s r(480)=0.66, p=3*10^−61^; monkey 2: Pearson’s r(480)=0.47, p=3*10^−27^). The strength of these correlations is notable given that it rivals the strength of correlations reflected in visual cortex for visually-evoked responses (41) but in the case presented here for AIC, this happens during the behaviorally unconstrained intertrial interval period and the variable in question is based on subjective happiness reports.

Qualitatively similar results were obtained with the CRTR model (Supplementary Fig 5A-I), including that correlation with model terms reflected an integration across ∼6 trials and they had a similar pattern across correlations with behavioral happiness reports versus AIC neural data (compare Fig 1F and Supplementary Fig 5H).

We wondered whether mood might be reflected not just in AIC but ubiquitously across the many higher brain areas that have been shown to reflect persistent activity. For insight, we applied the same CRTE model-based analyses to units recorded in lateral prefrontal cortex (LPFC) and there we found a very different result. Only 6.3% of LPFC units were significantly correlated with estimated happiness (significance criterion: p<0.001; Fig 3J; compared with 41% in AIC, Fig 3G). Among significant units, correlation magnitudes were weaker in LPFC than in AIC (Fig 3K; compare with Fig 3H). When LPFC units were configured into a pseudopopulation (the responses of 224 units on 480 trials), estimated happiness was very weakly negatively and non-significantly correlated with the overall vigor of the LPFC population response (Fig 3L; Pearson’s r(480)=-0.05; p=0.27). Similar results were obtained with the CRTR model (Supplementary Fig 5J-L). These results suggest that the behavioral signatures of mood are reflected much more strongly in AIC than in LPFC and are thus not generalized across all brain areas with persistent activity.

As mentioned above, the CRTE model was one of a class of several behavioral models we considered (Supplementary Table 2). The CRTE model estimated happiness based on 6 parameters that captured concrete rewards and token-related events parsed into 3 terms (model 6.1, concrete and token rewards, mean r^2^ = 0.38). In comparison, the worst-predicting behavioral model estimated happiness based on the total token number alone (model 3.3, mean Pearson’s r^2^=0.09). Variation in the average goodness of prediction that exists between these models (mean Pearson’s r^2^=0.09 - 0.38) provides an opportunity to assess whether the factors that make for a better versus worse model of human subjective happiness are related to the factors that make for a better versus worse model of what drives the vigor of the persistent firing rates in AIC. Indeed, behavioral predictions had a strong and significant relationship with AIC neural predictions in which rank-order was largely preserved (Figure 4, Pearson’s r(10)=0.82; p=0.004). These results are consistent with the hypothesis that human subjective happiness reports are reflected in monkey AIC as persistent activity.

**Figure 4.**
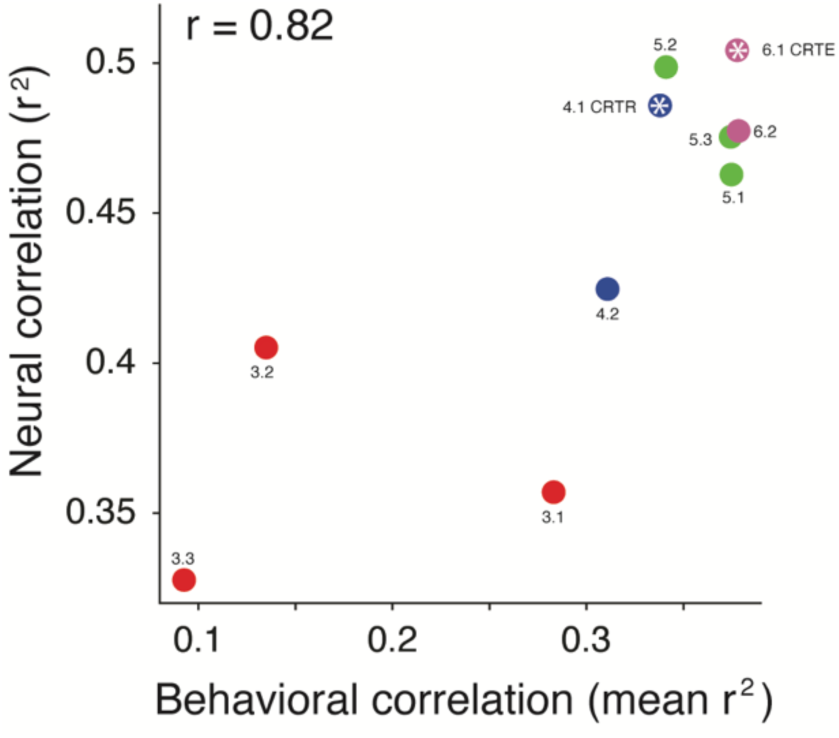
*Model comparison for behavioral versus neural data.* Plotted are the mean r^2^ for each behavioral model (Supplementary Table 2) versus the correlation of each model with AIC neural data (e.g. Fig 3I) with each model labeled. The CRTR model (4.1) and CRTE model (6.1) used in Fig 1 are labeled with asterisks. Colors reflect the number of parameters in each model (3: red; 4: blue; 5: green; 6: magenta).

As a final set of analyses on this data, we used the suite of mood models to investigate whether persistent activity in AIC reflected memories of past rewards as compared to the anticipation of future ones. Because the gambling paradigm involves the accumulation of tokens to receive concrete rewards, the two are correlated in this experiment (e.g. following a trial in which 3 tokens are won, persistent activity could reflect the token reward on the last trial or the anticipation that concrete reward may be received on the next one). However, following trials in which a concrete reward is delivered, 6 tokens are subtracted, and consequently, the probability of receiving a reward on the next trial is zero (because a concrete reward requires 6 tokens and the maximum that can be won is 3). As such, the ability of the concrete reward model alone (model 3.1) to account for the behavioral (Pearson’s r^2^ = 0.28) and neural (Pearson’s r^2^ = 0.36) data cannot be explained by reward anticipation; it can only be explained by reward memory. In comparison, a model based on reward anticipation alone loosely maps onto total token number (model 3.3), and of all the models we considered, this model provided the worst account of our behavioral (Pearson’s r^2^=0.09) and neural (Pearson’s r^2^=0.33) data. Thus, while we cannot rule out the possibility that reward anticipation contributes in some fashion to persistent activity in AIC, we can conclude it is not the primary driving force behind it.

Lastly, our results fall into intriguing alignment with a study reporting that average reward history across multiple trials impacts monkeys’ choices in a probabilistic reward learning task, where a component of neural activity in a similar region of monkey AIC measured by fMRI tracked average reward state (42). In that study, the authors did not link reward history to mood but instead to an integrative process used to drive reinforcement learning, similar to other reinforcement learning models that consider running average reward rate (43–45). This parallel leads to the question: does the persistent representation of reward history identified in AIC drive the subjective experience of mood, a running average of reward history for reinforcement learning, or both? Given the correlative nature of their study and our own, we cannot be certain, but we suspect the answer is “both”. This notion is central to theories that account for the relationship between decision-making and affect, including Decision Affect theory (46) and Integrated Advantage theory (1). The gist behind the latter is that a running average of Advantage (which in this task translates approximately to a running average of reward; Supplementary Fig 6) is computed for strategic purposes to drive reinforcement learning, and this is what humans perceive as mood. This proposal leads to the prediction that when humans are engaged in the task reported in (42) for monkeys, their subjective happiness will fluctuate with the running average of reward history. We tested this prediction in 55 human subjects, and it was confirmed (Supplementary Fig 6).

## Discussion

Because mood is a subjective experience that we have no external way to measure, studying how the brain supports it requires different approaches than studying many other brain functions (like perception, memory, and decision making). Here, we took a nontraditional approach to the challenge of studying mood by creating subject-generalized models to predict how the happiness of humans will fluctuate as they engage in a gambling task (based on the wins and losses they experience), coupled with investigation of the neural correlates of these fluctuations in the monkey brain.

We demonstrate a remarkably strong alignment between estimated happiness, monkey choice behavior, and the responses of units in AIC. Within AIC, 38% of neurons were significantly modulated by estimated happiness, and estimated happiness was strongly correlated with an estimate of population response vigor (r=0.71). The strength of this alignment is notable given that it rivals that of visual cortex for image properties that also modulate the population response (41). Of note is that in AIC, this alignment happens during the behaviorally unconstrained intertrial interval period (as opposed to a visually-evoked response) and reflects a behavioral index that is a proxy for subjective mood/happiness. In addition, we found that both happiness reports (Fig 1H) and the firing rate of AIC units (Fig 3H) correlated with reward-related events across similar numbers of trials (∼7). Furthermore, the CRTE model was significantly better than the CRTR model at predicting subject-left out human happiness reports (Fig 1G vs J) and monkey choice (Fig 2), and it was more correlated with AIC activation (Fig 4, compare blue and magenta dots with asterisks). Likewise, the factors that made for better versus worse behavioral models were correlated with those that accounted for AIC response vigor (Fig 4). In comparison, estimated happiness was only weakly reflected in LPFC (Fig 3J-L), suggesting that our results do not follow from a signal broadcast across all higher brain areas. Together, these results suggest that persistent activity in AIC may be a neural correlate of the subjective experience of human happiness in the context of this gambling task. Even if this is not true (because the neural correlate lies elsewhere or involves AIC only in part), these results shed insight into how happiness is computed in the primate brain.

Namely, our results inform how the brain transforms positive and negative experiences into mood during the gambling mood induction paradigm, constrained by the following signatures. First, mood is reflected in AIC as persistent activity. Second, mood depends on both concrete and token rewards, with token rewards parsed into quantities reflecting outcomes when certain versus gamble options were selected. Third, the computation of mood integrates both types of events across the past ∼7 trials, as a fading average. Fourth, in AIC the neural correlates of mood are dominated by neurons that positively correlate with happiness, but negatively correlated neurons exist as well (with ∼3.8x more positively tuned units).

Together, these results suggest a network model that performs integration of the relevant reward-related events and creates persistent activity by operating via recurrently connected push-pull pools of positive and negative neurons (Fig 5) (47). Here, we extend those ideas to sketch a model of how mood is computed in the brain — which may happen within AIC or may happen elsewhere and simply be reflected there. In this framework, mood is computed based on experiences of concrete and token rewards which are fed into a recurrent network configured to integrate information across ∼7 trials via reciprocally connected pools of excitatory (E) and inhibitory (I) units (Figure 5). The resulting network output then reflects mood as persistent activity, including population response vigor in AIC. The proposal in Fig 5 raises the intriguing possibility that individual proclivities to mood disorders may follow from individual differences in the biophysics and connectivity of this type of network.

**Figure 5.**
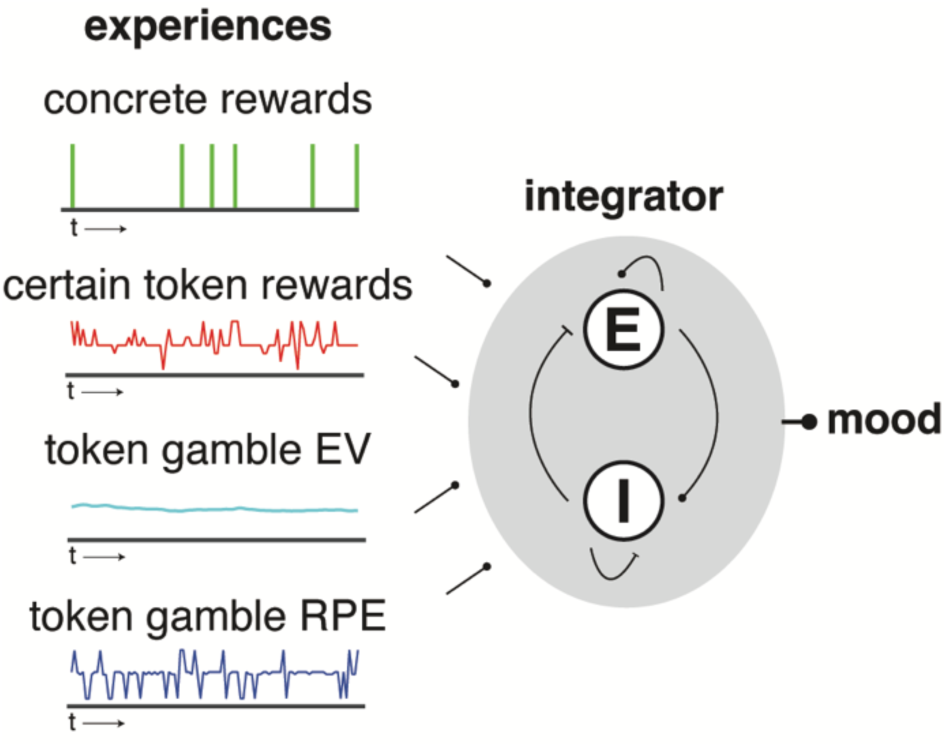
*An emerging sketch of how the brain transforms experiences into moods.* The fact that happiness reflects an integration of experiences, combined with the finding that it is reflected in AIC as persistent activity, is consistent with models in which integration happens via a population of reciprocally coupled excitatory and inhibitory neurons (47) that integrate reward-related input. In the CRTE model, those include *concrete rewards.* They also include token rewards parsed into *certain token rewards* (tokens acquired when the certain option was selected), *token gamble EV* (when the gamble option was selected, expected value computed as the average outcome on gamble trials up to the trial in question) and *token gamble RPE* (when the gamble option was selected, reward prediction error, the difference between expected value and the outcome).

Several other lines of evidence implicate AIC in the conscious experience of feelings including mood (36), including natural human lesions (48), fMRI (29, 35, 49), and microstimulation (50, 51). Prior work in humans has also demonstrated correlations via fMRI (29) and broadband gamma activity (52) (often considered a proxy for average spiking activity) in AIC with human subjective reports of mood following feedback on a trivia quiz. Our study complements this other work by providing insight into the nature of mood representations in AIC at the resolution of individual units and their action potentials. Like the bulk of neurophysiology studies, the results we present here are correlative. Based upon them, we do not know whether persistent activity in AIC drives happiness or even if it is necessary for it. However, these results lay the foundation required to design causal perturbation experiments to investigate that question.

Our recordings largely targeted dorsal AIC, and within the region we studied, we found no obvious anatomical organization of the units that significantly reflected estimated happiness versus ones that did not (Supplementary Fig 7). Across a larger region, the insula is organized anatomically and functionally along both its anterior-posterior and dorsal-ventral axes (reviewed by (36, 53, 54)). Currently, the anatomical components of this organization are best understood in macaque monkeys, whereas the functional organization is best understood in humans; the functional mapping between human and macaque insula is incompletely understood (55). In humans, meta analyses tie emotion-related functions to AIC with social-emotional functions tied to its ventral region and cognitive functions tied to its dorsal region (56). Intriguingly, one study found that stimulation of human AIC during a risky decision-making task impacted loss aversion in a manner that depended on the region of AIC being stimulated, where stimulation of dorsal AIC decreased loss aversion, and stimulation of ventral AIC increased it (57). The former aligns with our findings that happiness decreases loss aversion in monkeys (Fig 2, red dotted versus solid lines) and it is reflected as population response vigor in dorsal AIC, under the assumption that stimulation increases population vigor. Of great interest but still unknown is whether stimulation of human dorsal AIC impacts the subjective experience of happiness.

While running average reward rate integrated across concrete and token rewards (the CRTR model) was not the best among all the models we considered for human subjective happiness, monkey choice modulation or AIC neural responses, it provided a reasonable estimate of all three. Another study has also linked AIC neural activity (measured in that case by fMRI) with running average reward history (42). In that study, the link was made in the context of a probabilistic reward learning task and the interpretation was that integrated reward likely exists in AIC for strategic purposes. Consistent with a theory that links reward history in this context with the subjective experience of mood (1), we predicted that when humans engaged in the same task as reported in that study, subjective happiness would also fluctuate with running average reward history — and this prediction was confirmed (Supplementary Fig 6). As such, evidence now exists across two paradigms linking quantities approximated by running average reward history with an influence on monkey choice during risky decision making, neural activity in monkey AIC, and human subjective experience. A parsimonious account of both studies is thus that quantities related to running average reward history are computed by the primate brain for strategic purposes, and this is what humans perceive as mood.

Here we report a strong link between AIC activity and mood, but the network generating and representing mood is likely more wide-spread. Attempts to decode mood from the human brain via intracranial electrocorticogram (ECoG) measures have implicated a host of other limbic brain areas outside of AIC in mood (such as cingulate and orbitofrontal cortex as well as the amygdala and hippocampus, (16, 17)). Of note, other studies have found evidence for persistent representations of integrated reward in some other brain areas, in particular anterior cingulate cortex (ACC, (58)) where (as in our study of AIC), the fraction of units of this type was significantly higher than LPFC (59). These results are consistent with strong functional homologies between AIC and ACC, although AIC has been more strongly associated with emotional awareness (60). In comparison, persistent representations of reward in ACC have been linked to the ongoing evaluation of choice outcomes based on reward history in a game that requires continuous monitoring of strategy and adjustments of it (58). These two interpretations of the function for reward-related persistent activity are not mutually exclusive, as happiness not only reflects integrated experience but in turn, also impacts decision making, as shown here (Figure 2) and in previous findings (1, 21, 29, 31–34, 38). Analogous persistent representations of reward across trials have also been demonstrated in rodents (61–63).

The gambling mood induction paradigm we employ here has yet to be fully characterized in terms of modulations of valence and arousal in ways analogous to other approaches (e.g. (64)). Likewise, future work will be required to determine how these results relate to mood when it is modulated by other induction paradigms such as those that use audiovisual stimuli (65) or autobiographical recollection (66). These different approaches to mood have different strengths and drawbacks. The strength of the gambling paradigm we use here is that it can easily be quantified and incorporated into models of what subjective happiness depends upon (Fig 1). While foundational steps toward predicting affective states from images and videos are beginning to emerge (67), comparable approaches for audiovisual stimuli and autobiographical memory have not been developed. As such, there is currently no replacement in those domains that allows for the type of model-based, cross-species comparison we perform here.

Finally, while we have no way of determining what the monkeys experience as they perform this task, our approach is grounded in the subjective experience of humans, which may be essential for arriving at an impactful understanding of mood. For emotions such as fear, concerns have been raised that the circuits that mediate innate behaviors (including threat-related behaviors such as freezing) may be distinct from those that mediate subjective experience and consequently, the behaviors that mediate innate behavior may be of limited therapeutic relevance (14). By focusing on the neural correlates of human subjective experience, our approach seeks to bridge between neuroscience work in humans and animals using a model-based approach. Going forward, a parallel pursuit of all three approaches — work in humans, work in animals, and approaches that bridge across species— will be crucial for an impactful understanding of mood.

## Methods

As an overview, data from two sources are included in this paper. First, behavioral and neural data collected from monkeys performing a gambling task — including a study also reported here (37). That report examined the relationship between neural responses in anterior insular cortex (AIC) and decisions in a gambling task to test ideas related to the prospect theory of decision making. The results presented here focus exclusively on an epoch that was not analyzed in that report (the period between trials) and these results cannot be inferred from that paper. We also include unpublished data collected from the same two monkeys in the original report, but from lateral prefrontal cortex (LPFC).

In addition, this study includes behavioral data collected from humans performing the same gambling task with additional subjective happiness probes, inspired by the approaches reported by (35), with modifications. We also include behavioral data collected from humans performing a probabilistic reward learning task linked to AIC (42), also with happiness probes inserted.

The Methods associated with each type of data are described below. Upon acceptance (and finalization of the code), all data used for the paper and analysis code will be placed in a publicly accessible repository.

### Behavioral and neural data collected from monkeys performing a gambling task (absent happiness probes)

Because the methods used to collect this data are described in detail by (37), we summarize them here. All animal care and experimental procedures were conducted in accordance with the US public Health Service policy on the humane care and use of laboratory animals and were approved by the Johns Hopkins University Institutional Animal Care and Use Committee (IACUC).

Experiments were performed on two adult male rhesus macaque monkeys (Monkey 1: 7.2 kg, 7 years; Monkey 2: 9.5 kg, 8 years). All behavioral training and testing were performed using standard operant conditioning (juice reward), head stabilization, and high-accuracy, infrared video eye tracking using MonkeyLogic software (68). (https://www.brown.edu/Research/monkeylogic/) was used to control task events, stimuli, and reward, as well as monitor and store behavioral events.

We used T1 and T2 magnetic resonance images (MRIs) (3.0 T; Johns Hopkins Hospital) to determine the recording locations in both the anterior insular cortex (AIC) and lateral prefrontal cortex (LPFC) of each monkey. AIC, situated within the Sylvian (lateral) fissure, is implicated in interoception, emotion, and mood regulation. It is bordered anteriorly by the orbital prefrontal cortex and covered dorsally and ventrally by the frontoparietal and temporal opercula, respectively. LPFC, located in the frontal lobe and involved in cognitive control and decision-making, was targeted near the principal sulcus, specifically in areas 8Ad and 8Av, slightly dorsal to the posterior principal sulcus. Recording sites in AIC and LPFC were estimated using the stereotaxic coordinates of the recording chamber and electrode penetration depths, and superimposed onto MRI scans (Supplementary Figures 7 and 8).

Single neuron activities were recorded extracellularly with single tungsten microelectrodes using the TDT system (Tucker & Davis). Action potentials were amplified, filtered, and discriminated conventionally with a time–amplitude window discriminator. Neural data were recorded from anterior insular cortex, guided by stereoscopic chamber placement based on MRI images, and recording electrode depth.

AIC neural data include 235 unique units, parsed into 137 neurons recorded in monkey 1 and 98 neurons recorded in monkey 2. LPFC data included 224 unique units, parsed into 144 units in monkey 1 and 100 in monkey 2. Both data sets were collected as the animals performed the gambling task.

To compare units (e.g. Fig 3G-L), we placed a threshold on numbers of trials (>479) and equated their numbers of trials to 480. This resulted in a modest (<4%) reduction in the numbers of units from 235 to 229 in AIC and no reduction in LPFC. To ensure that subsampling did not impact the distributions of sampled estimated happiness scores, we ranked trials by their estimated happiness and subsampled them evenly to achieve 480. For instance, for a unit with 480*2 trials, every second trial would be kept after ranking by estimated happiness.

### Behavioral data collected from humans performing gambling tasks (with happiness probes)

International, English-speaking subjects (n=55 for both behavioral studies) were recruited through the online website Prolific and provided informed consent to participate. The two tasks were run on the online platform Pavlovia driven by PsychoPy code. Subjects were adults (Study 1: age range 18-63, median 33; Study 2: age range 19-78, median 39) that self-identified as Study 1: 47% male, 48% female, 5% undisclosed and Study 2: 60% male, 40% female, 0% undisclosed. Human behavioral experiments were conducted with the approval of the University of Pennsylvania Institutional Review Board for the Protection of Human Subjects (IRB).

### The token-based gambling task

The token-based gambling task consisted of two types of trials: choice trials and forced choice trials. In choice trials, two targets (a sure option and a gamble option) were presented on the screen. In forced choice trials, a single target was presented. In both cases, the token values of the target(s) were indicated. For monkeys, the token values corresponding to options were indicated by colored boxes; for humans, numbers were displayed (Figure 1). Monkeys selected an option by making a saccade to it; humans selected an option with a button press.

Each trial began with a display of the token count, followed by the options and a selection. When the total token count was greater than 6, a reward was delivered at the end of the trial (a liquid reward for monkeys; a monetary reward of $0.50 for humans) and 6 tokens were subtracted from the total. When the token count fell below 0, it was set to zero. Humans began the experiment with a $6.00 base reward.

Here, we focus primarily on the neural representation in the insula during the period between trials, which always exceeded 1.7 seconds. During this period, the monkeys’ behavior, including their eye movements and location of focus, was unconstrained.

The total number of choice conditions was 24, including all possible combinations of 4 sure options and 3 gamble options (for a total of 12) in symmetric gain (win token) and loss (lose token) conditions (for a total of 24). In addition, each of the 13 options were presented alone on forced choice trials. The 37 trial types were presented in a pseudorandom block design in which each trial type was presented before re-randomizing the sequence. In the human experiments, 4 blocks in total were presented. In the monkeys, the number varied from session-to-session; the minimum in this dataset was 12.

For human subjects, a happiness probe was interleaved every 2-3 trials. On these trials, subjects were presented with the question, “How happy are you at this moment (1–20)?” Subjects entered their rating (from 1-20) with a button press. Subjects completed a practice session before data were collected.

### The probabilistic reward learning task

The probabilistic reward learning task mimicked the design of (42). It included 3 pictures, each associated with a different reward probability trajectory across the 200 trials of the experiment (Supplementary Fig 6a). For two pictures, the average probability flipped halfway through the experiment; for the third picture, reward probability was sinusoidal across the 200 trials. Humans were instructed, “There are three total images you might see during the experiment. Each image is associated with a different reward probability. For some images, those probabilities will change part way through the experiment. Your goal is to infer the underlying probabilities to help guide your choices such that you maximize your reward”.

On each trial, subjects were presented with two of the three pictures and selected one; whether $0.10 was awarded or not was computed according to the probability of receiving a reward for that picture on that trial. Happiness probes were inserted every 2-3 trials. Humans began the experiment with a $5.00 base reward.

### Subject-generalized mood model

For the token-based gambling task, we compared a number of models to determine the ones that provided the best accounts of the behavioral data (Supplementary Tables 1-2). Following on Rutledge et al., every model had two parameters in common: a baseline happiness term *w_0_* and a parameter that determines the number of trials to integrate over (*γ)* in the equation below. In addition, each model had one or more parameters that corresponded to the weights w_n_ for each variable included in the model. The generic form of the model can be described as:

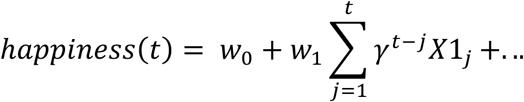

where w_1_ is the weight for the variable X1. For instance, the CRTR model used in Fig 1D-G had 4 parameters (*w_0_-w_2_*, and *γ*):

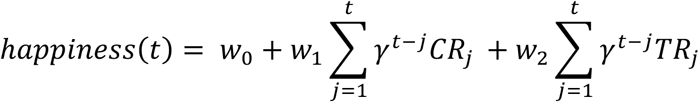

Here, TR refers to the number of tokens won on each trial (token reward) and CR to whether a concrete reward was received on that trial (which happened when tokens reached 6).

The parameters were fit to the combined data across 54 human subject using the optimization toolbox in MATLAB. They were then used to estimate happiness for the each of the 55th subject’s held-out data. To predict happiness fluctuations during the experiment (as opposed to the factors, possibly external, that contributed to the baseline), happiness measures for each subject were z-scored and the *w_0_* term was not used to make model predictions. To estimate happiness for the monkeys, we fit the subject-generalized mood model to all 55 human subjects’ data.

### Model of monkey choice behavior

We modeled the probability that monkeys chose the gamble option by a softmax choice function whose argument was the difference between the expected utility of each option. Utility was parameterized as:

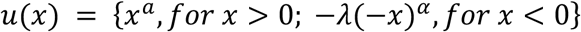

where α is a parameter determining the curvature of the utility function, u(x), and *x* is the reward outcome (in units of gaining or losing token numbers). λ indicates the modulation of utility function in loss context. α and λ were computed as a weighted sum of a constant (α_0_ and λ_0_), one term for token asset modulation (α_T_ and λ_T_), and a second term for happiness modulation (α_H_ and λ_H_):

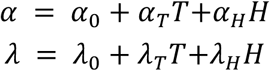

where T reflects current token assets and H estimated happiness on each trial. Model performance was compared with a second model in which α and λ were computed absent the happiness term:

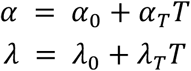

Subjective probability of each option was computed by:

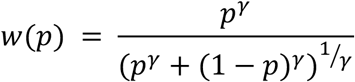

where γ is a free parameter determining the curvature of the probability weighting function, *w(p)*, and *p* is the objective probability of receiving the corresponding outcome. The expected utility (EU) of each option was computed by combining the output of *u(x)*; and *w(p)* that map objective gains and losses relative to the reference point and objective probability onto subjective quantities, respectively:

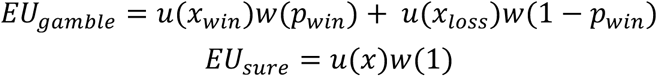

The expected utility difference between the two options was then transformed into choice probabilities via a softmax function with terms of slope *s* and bias *s*:

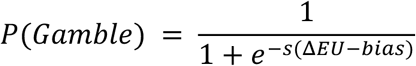

where *ΔEU = EU_gamble_ - EU_sure_*, *s* determines the sensitivity of choices to the ΔEU, and bias is the directional bias of choosing gamble.

We optimized the 9 model parameters, α_0_, α_T_, α_H_, λ_0_, λ_T_, λ_H_ γ, *s* and *bias* in the full model (with 2 happiness terms), separately when happiness was estimated with the CRTE and CRTR models. We also optimized a 7 parameter model with no happiness terms with parameters α_0_, α_T_, λ_0_, λ_T_, γ, *s* and *bias*. Models were optimized by minimizing the negative log likelihoods of the data given different parameter settings using Matlab’s fmincon function, initialized at multiple starting points of the parameter space as follows:

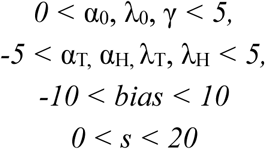

Negative log-likelihoods (-LLmax, which measures the accuracy of the fit) was used to compute classical model selection criteria. We compared models via one-sided paired t-tests to compare - LL_max_ calculated from a 5-fold cross-validation (with 20% of data held out each time), and these are reported in the Results. Complementary comparisons via the likelihood ratio test following model fits to all data are presented in Supplementary Table 3.

### Population-based analysis of neural data

Accurate estimate of population response magnitude requires many units to eliminate the selectivity (sometimes called “tuning”) of individual sampled units. To perform our analyses, we concatenated units across sessions to create a pseudopopulation using methods analogous to those described by (41). Specifically, when creating this pseudopopulation, we aligned data across sessions in a manner that preserved estimated happiness. The number of trials that could be included in the pseudopopulation was limited by the session for which the fewest trials were obtained across both monkeys. For the other sessions, a matched number of trials was subselected by ranking happiness for that session, preserving the lowest-ranked and highest-ranked happiness within that session, and selecting the number of additional images required as those with estimated happiness that were evenly spaced between the two extreme happiness estimates for that session.

The resulting pseudopopulation consisted of the responses on 480 trials. To perform the neural analyses (Fig 3J, Fig 3M, Fig 4), an estimated happiness for each of the 480 trials was computed as the mean of the estimated happiness scores that were aligned to produce the pseudopopulation response for that trial. Our analyses focus on the hypothesis that a relationship exists between the vigor of the population response and estimated happiness. As such, aligning the population in this way is not circular insofar as if such a relationship did not exist, our analysis would produce a null result.

## Acknowledgements

This work was supported by the Simons Foundation (Simons Collaboration on the Global Brain award 543033 to N.C.R., a Pivot Fellowship to N.C.R., and the National Science and Technology Council (NSTC, Taiwan) grant 113-2320-B-016 −002 to Y.Y.

## Supplementary information

**Supplemental Figure 1.**
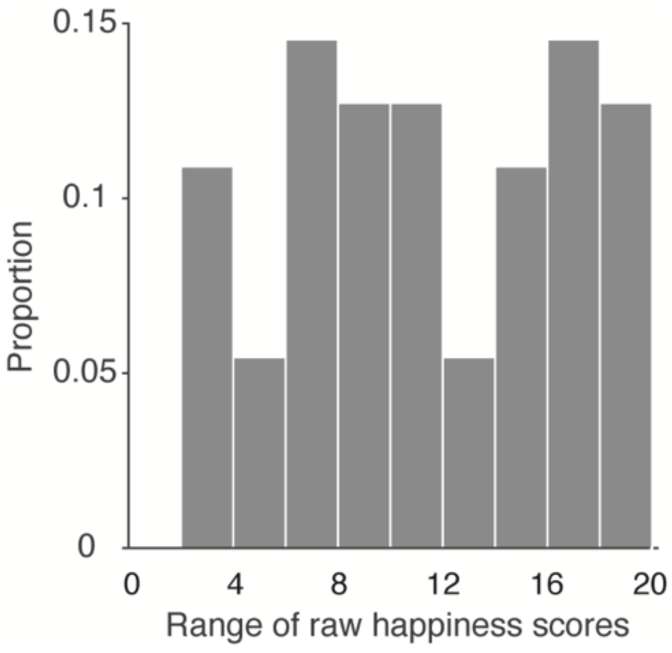
R*a*nges *of happiness scores used by individual subjects.* Here, range was computed as the difference between the highest and lowest happiness score used by each of 55 human subjects (who were asked to scale their subjective happiness 1-20). 100% of subjects used a range more than 2, 95% of subjects more than 4, and 84% of subjects more than 6. The mean and std range was 11.8 +/− 5.1.

**Supplemental Figure 2.**
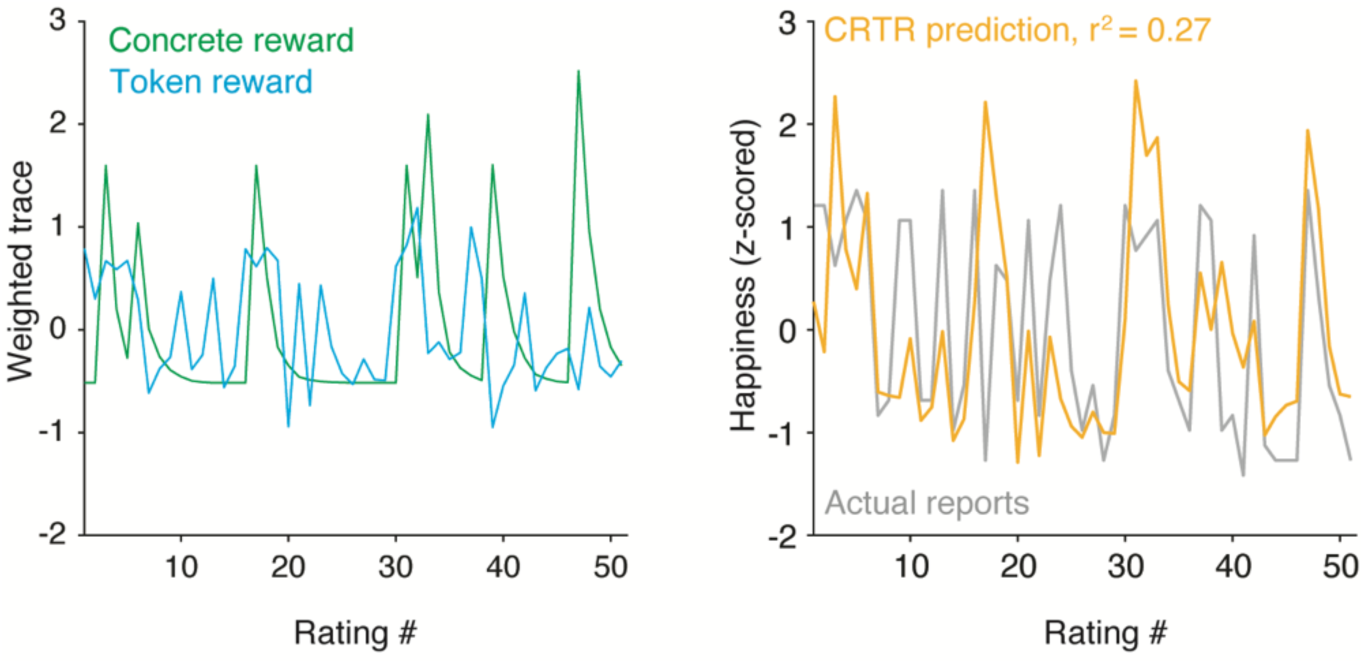
C*o*mputation *of CRTR happiness model predictions.* Right: Actual happiness fluctuations (gray) and CRTR happiness model predictions (yellow), shown for the same subject depicted in Fig 1D. Left: those model predictions were computed as a sum of fluctuations in monetary reward (green) and token number (cyan). Each type of fluctuation was computed by taking the events that happened during the course of the experiment (monetary and token rewards) and integrating them both across time and with one another by passing them through the weights shown in Fig 1F.

**Supplemental Figure 3.**
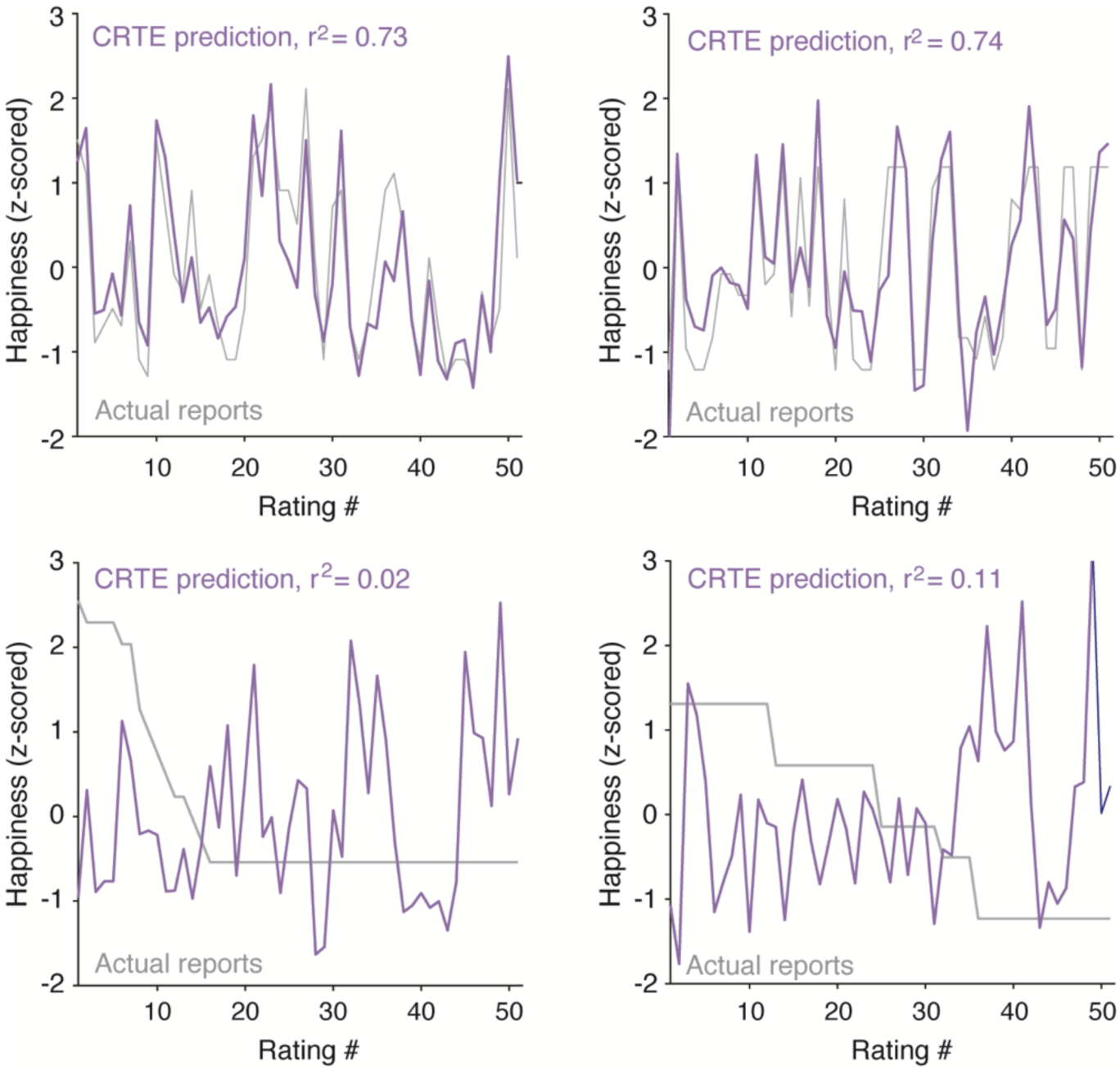
F*o*ur *additional example subjects. Top:* two subjects with exceptional CRTE model predictions. *Bottom:* two subjects whose subjective happiness reports were insensitive to the events that happened during the gambling mood induction paradigm. In these subjects, happiness decreased across the experiment (possibly due to boredom). Removing these subjects and the handful of others whose happiness reports were not strongly correlated with events during the gambling experiment did not qualitatively impact the results (not shown).

**Supplemental Figure 4.**
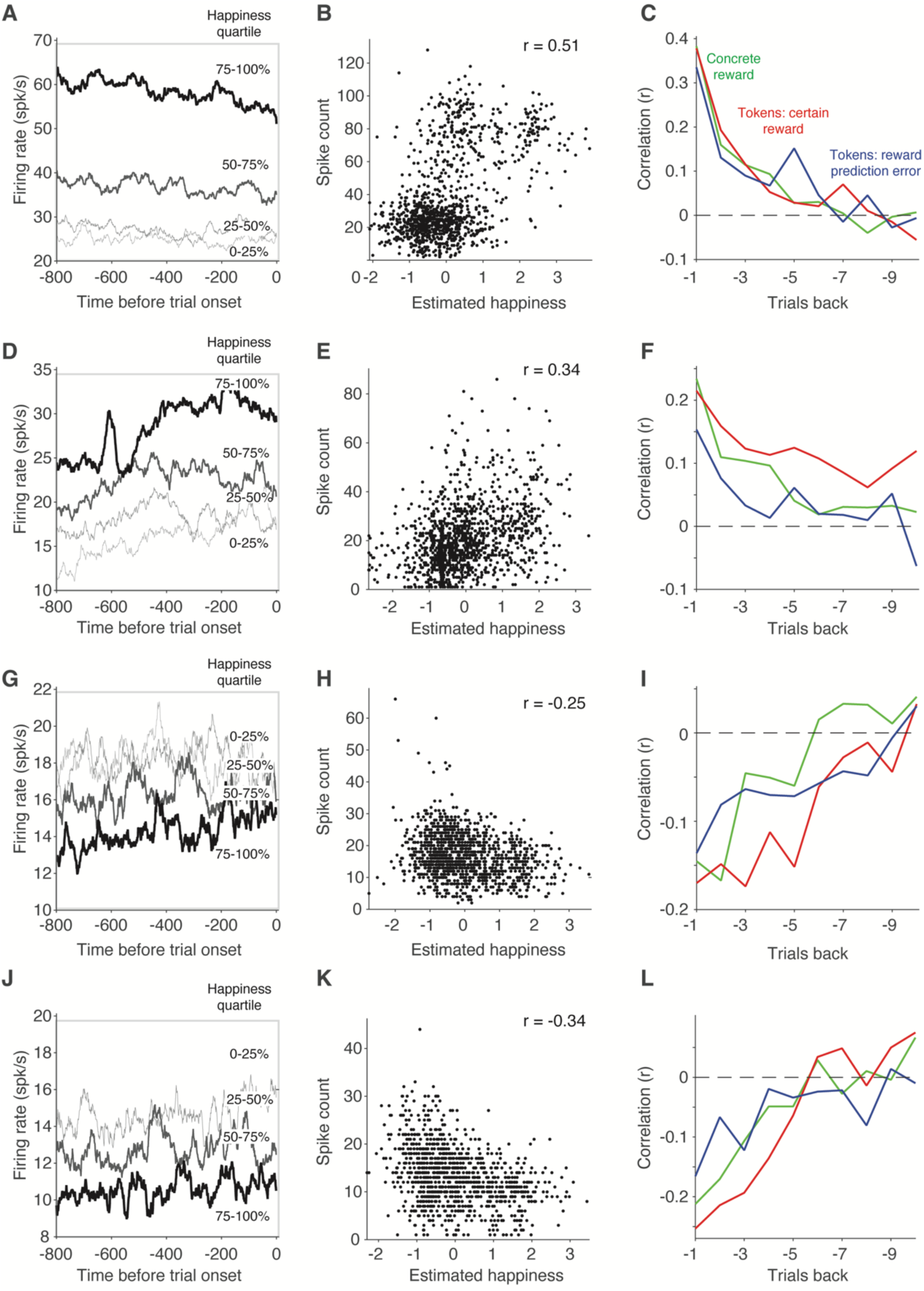
T*h*e *reflection of happiness as estimated by the CRTE model for four additional example units.* Examples were selected to depict a range of correlations. Shown are a positive (A-C) and negative (J-L) unit with strong correlations, and two units near the mean correlation across all neurons with significantly positive (D-F) and negative (G-I) correlations. Panels are plotted with the same conventions as Fig 2 A-F. Number of trials: A-C) 1181, D-F) 1257, G-I) 1254, J-L) 1060.

**Supplemental Figure 5.**
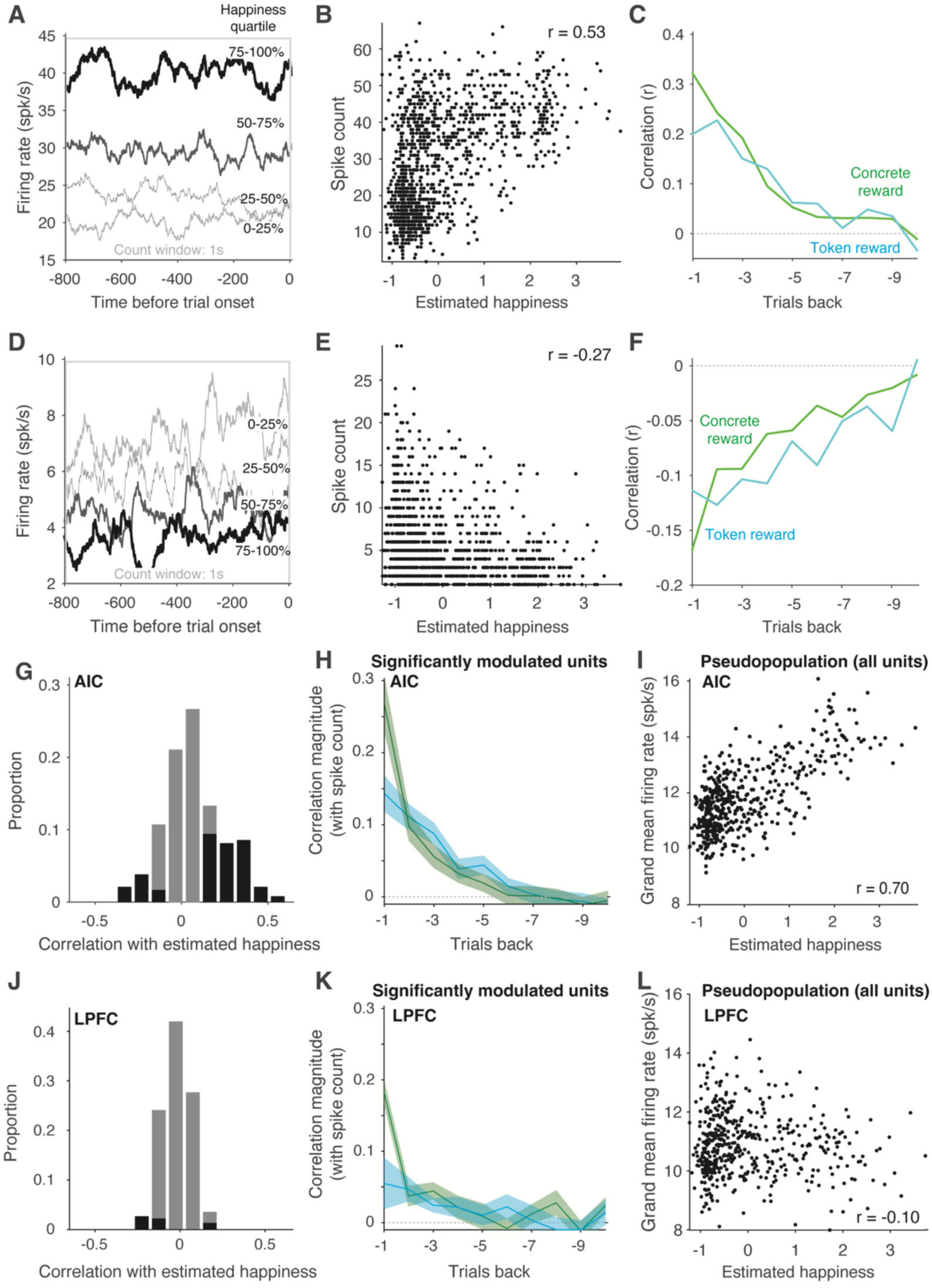
I*n*dividual *AIC units reflect happiness estimated by the CRTR model.* Shown for comparison with Fig 3 (where happiness was estimated with the CRTE model), with the same conventions. B) r(1307) = 0.53; p = 2*10^−94^. E) r(1276) = −0.27; p = 2*10^−23^. G) 38% of AIC neurons were significantly correlated with estimated happiness, with 3.8-fold more positive than negative; significance criterion p<0.001. I) r(480) = 0.70; p = 5*10^−71^. J) 6.3% of LPFC neurons were significantly correlated with happiness; significance criterion p<0.001. L) r(480) = −0.10; p = 0.03. For the analyses in G-L, units were limited to 480 trials via equal spacing across the sampled estimated happiness range to equate between them.

**Supplemental Figure 6.**
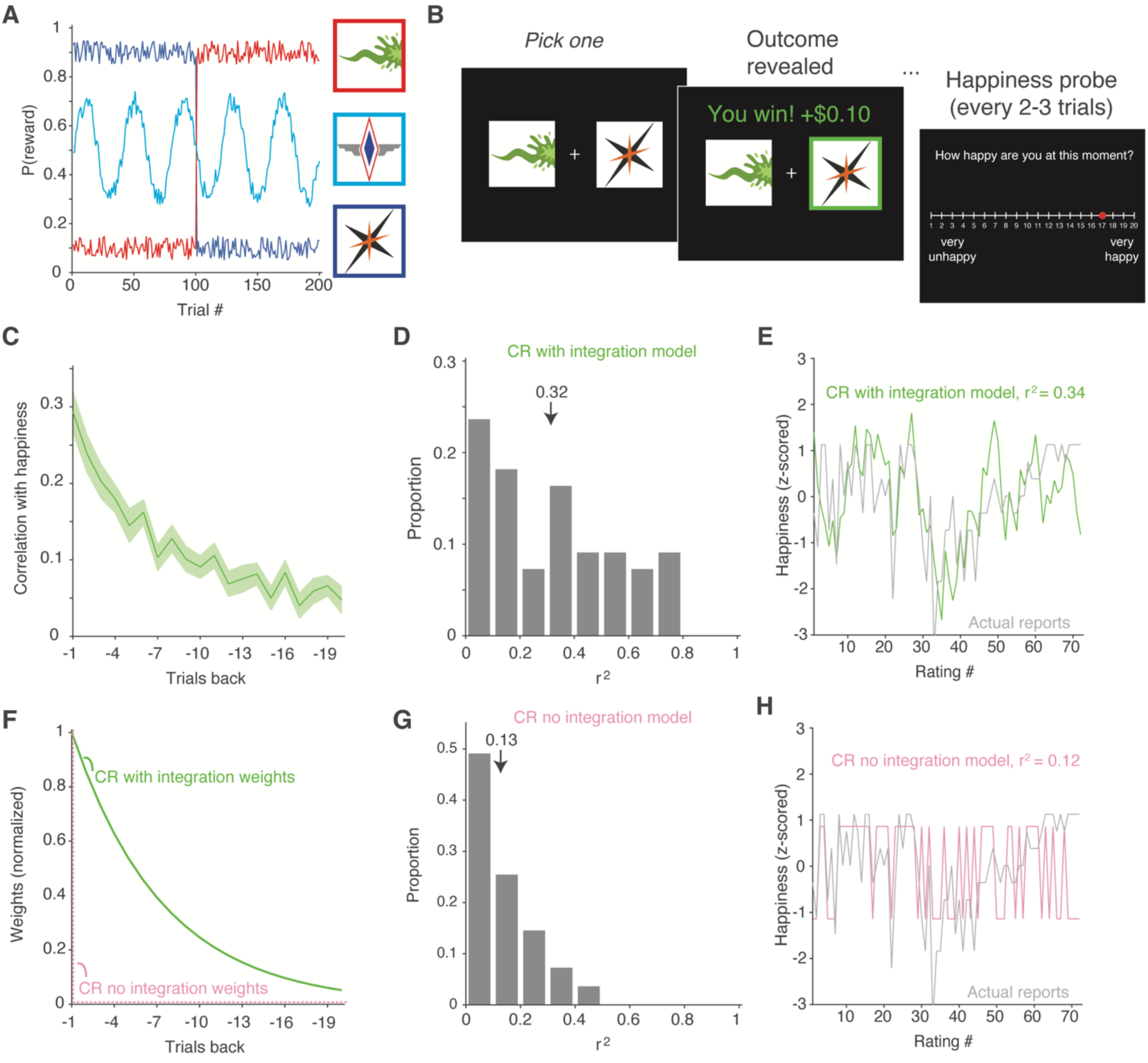
M*o*od *model for a second task linking AIC neural responses and integrated reward history.* In a study reported by (1), integrated reward history was linked to monkey choice and monkey AIC activity measured by fMRI, not in the context of mood but a probabilistic reward learning task. Here we test the hypothesis that during that task, human subjective happiness fluctuates with integrated reward history. This prediction follows from the alignment between the equation used by (1) to model reward history (which they called R_trace_) and the equation for Advantage specified by (2) (see also the mood equation in (3)). While Advantage is computed based on prediction error (the difference between actual and expected rewards), expected rewards (computed as the average of all rewards up to the trial in question) quickly converges to a constant value in this paradigm after the first few trials and after the subjects adjust their behavior following a switch, leading to the approximate equivalency. **A)** The experiment in (1) included 3 pictures, each associated with a different reward probability trajectory across the 200 trials of the experiment. For pictures 1 & 3, the average probability flipped halfway through the experiment. For picture 2, reward probability was sinusoidal across the 200 trials. Shown are the probability trajectories and pictures used in the human behavioral replication. **B)** On each trial of (1), the monkeys were presented with two of the three pictures and selected one; whether juice was awarded or not was computed according to the probability of receiving a reward for that picture on that trial (approximated by the traces in panel a). In the human replication, 55 humans performed the task described in (1) with happiness probes inserted every 2-3 trials. In lieu of juice rewards, humans received $0.10 added to their total when they won. Given the different nature of the task relative to Fig. 1 (where here there are only concrete rewards; no tokens), we focused on fitting a model in which happiness depended only on integrated concrete reward history (the “CR model with integration”) described as model 3.1 in Supplementary Table 2. We compared this with a reduced version of the same model where happiness depended only on the events on the last trial (“CR model with no integration”). **C)** Correlation between human happiness reports and concrete rewards, demonstrating the influence of both on happiness reaching back several trials. Lines indicate means and the shaded region indicates standard error across 55 subjects. **D)** Distribution of r^2^ for the CR model with integration across all 55 subjects determined by leaving out each subject in turn and fitting the model to the remaining 54 subjects’ data. Arrow indicates the mean. **E)** The actual and predicted happiness fluctuations for one representative subject. Shown are the predictions of the CR model with integration after fitting each model to the other 54 subjects’ data. **F)** The weighting functions for concrete reward with integration model following fits to all 55 subjects’ data combined (compare with panel C), along with the weighting function for concrete reward with no integration for comparison. **G)** Distribution of r^2^ for the CR model with no integration across all 55 subjects determined by leaving out each subject in turn and fitting the model to the remaining 54 subjects’ data. Mean r^2^ was significantly higher for the CR model with integration than without integration (Wilcoxon signed rank test, zval=5.76, p = 8.61*10^−9^). **H)** The actual and predicted happiness fluctuations for the same subject shown in E for the CR model without integration.

**Supplemental Figure 7.**
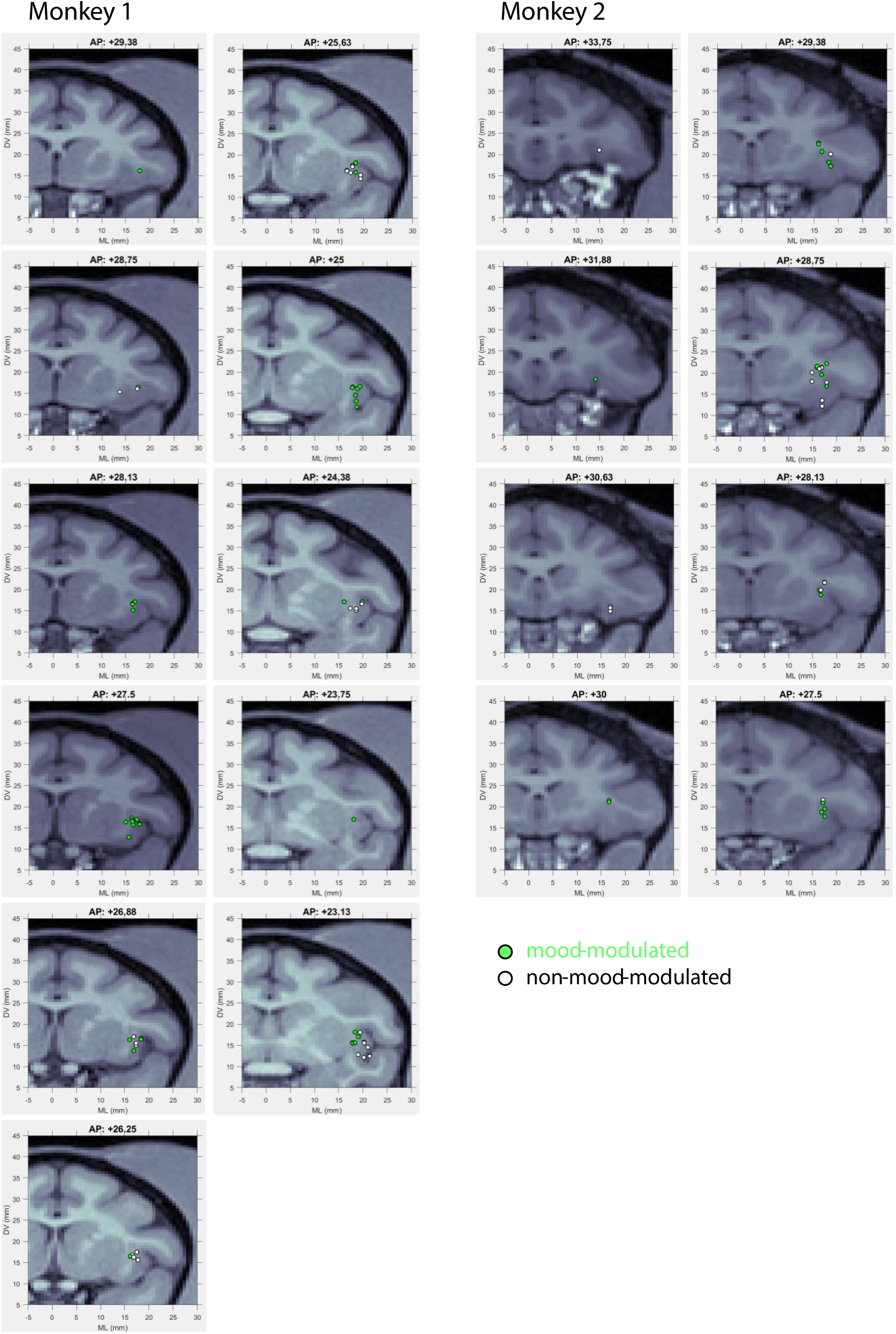
A*I*C *recording locations for Monkey 1 and Monkey 2.* Green indicates units that were significantly correlated with estimated happiness (Fig 3G).

**Supplemental Figure 8.**
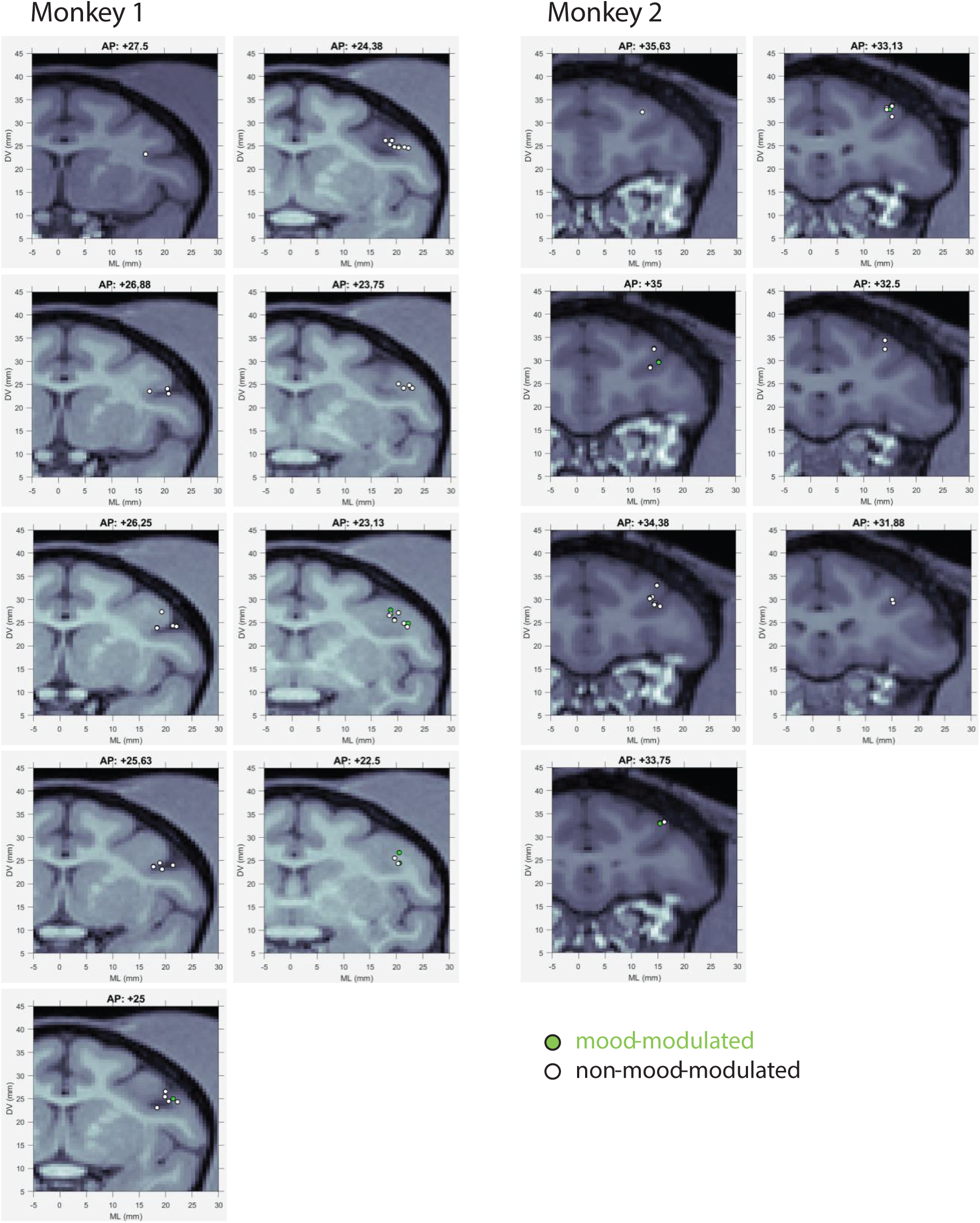
L*P*FC *recording locations for Monkey 1 and Monkey 2.* Green indicates units that were significantly correlated with estimated happiness (Fig 3J).

### Comparison of human behavioral mood models

In the study that inspired the behavioral component of this work, Rutledge et al. (2) compared the ability of a class of models to predict individual subjects’ happiness fluctuations as they won and lost money on each trial of a gambling task. In comparison, the gambling task used to collect the neural component of this work (Yang et al. (3)) differed in a notable way: subjects played for abstract, token rewards and received concrete ones (money for humans; juice for monkeys) when the number of tokens reached six. Here we thus must revisit the question of which model within the general class Rutledge et al. (2) considered accounts for happiness fluctuations with these modifications.

Our goal is also a bit different than the original Rutledge study insofar as we were focused on creating a model that captured subject-generalized fluctuations in happiness (as opposed to individual differences). To achieve this, we fit data combined across 54 subjects to predict happiness fluctuations for each of the 55th subject’s data. To predict happiness fluctuations during the experiment (as opposed to the factors, possibly external, that contributed to the baseline), happiness measures for each subject were z-scored and the *w_0_* term (described below) was not used to make model predictions. We measure goodness of prediction as the Pearson r^2^ between actual reports and held out model predictions.

Below we present these models by introducing their simplest versions first and then building them up to incorporate additional parameters. Following on Rutledge et al., every model has two parameters in common: a baseline happiness term *w_0_* and a parameter that determines the number of trials to integrate over (*γ)* in the equation below. In addition, each model had one or more parameters that corresponded to the weights w_n_ for each variable included in the model. For instance, the 3 parameter model that computed happiness based only on concrete reward (CR) took the form:

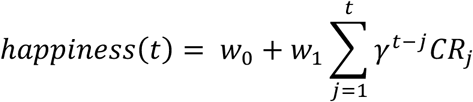

Similarly, the 4 parameter model that computed happiness based on concrete rewards (CR) and token rewards (TR) took the form:

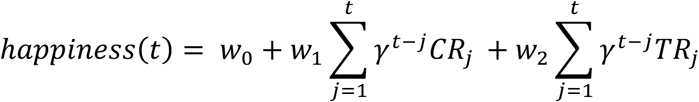

A description of all variables we considered can be found in Supplementary Table 1. The parameterization of all the models that we considered and their predictive power is summarized in Supplementary Table 2.

**Supplementary Table 1.** A description of all variables considered in the model fits.

|  |  |  |
| --- | --- | --- |
| CR | Concrete reward | Took on a value of 1 when a concrete reward was received and 0 otherwise. |
| TR | Token reward | Number of tokens won per trial; range -3 to 3 |
| TT | Total token number | Number of total tokens at trial end; range 0 to 6 |
| TS | Token start | Number of total tokens at trial start; range 0 to 5 |
| CrR | Certain rewards | Number of tokens won on trials when the certain reward option was selected (nan otherwise); range -3 to 3 |
| GR | Gamble rewards | Number of tokens won on trials when gamble option was selected (nan otherwise); range -3 to 3 |
| EV1 | Expected value (definition 1) | When gamble option was selected, probability weighted average between options (nan otherwise), as described by (2). |
| RPE1 | Reward prediction error (definition 1) | When gamble option was selected, difference between tokens won and EV1, as described by (2). |
| EV2 | Expected value (definition 2) | Average value of all token rewards received up to the trial in question (for both certain reward and gamble selections), as described by (4) |
| RPE2 | Reward prediction error (definition 2) | Difference between tokens won and EV2, (for both certain reward and gamble selections), as described by (4) |
| EV3 | Expected value (definition 3) | Average value of all token rewards received up to the trial in question, computed in the same manner as EV2 but only for gamble selections, as described by (4) |
| RPE3 | Reward prediction error (definition 3) | Difference between tokens won and EV3, only for gamble selections, as described by (4) |

**Supplementary Table 2.** Mean r^2^ between subjective happiness reports and model predictions, computed for 55 subjects. Models 4.1 (CRTR) and 6.1 (CRTE) were used in Fig 1; Model 6.1 (CRTE) was used to create figures 2-3. In Fig 4, all 10 models are compared.

| Number of parameters | Model name | Events | mean $r^2$<br>all subjects |
| --- | --- | --- | --- |
| 3 | 3.1 | CR | 0.28 |
| 3 | 3.2 | TR | 0.14 |
| 3 | 3.3 | TT | 0.09 |
| 4 | 4.1 (CRTR) | CR + TR | 0.34 |
| 4 | 4.2 | CR + TT | 0.31 |
| 5 | 5.1 | CR + CrR + GR | 0.38 |
| 5 | 5.2 | CR + TR + TS | 0.34 |
| 5 | 5.3 | CR + EV2 + RPE2 | 0.37 |
| 6 | 6.1 (CRTE) | CR + EV3 + RPE3 + CrR | 0.38 |
| 6 | 6.2 | CR + EV1 + RPE1 + CrR | 0.38 |

### Models based on single events (3 parameters)

The best fitting model of the single-event class computed happiness based on concrete rewards (CR; model 3.1, mean r^2^=0.28). Equivalent models based on token rewards (TR, model 3.2, mean r^2^=0.13) or total token number (TT, model 3.3, mean r^2^=0.09) provided worse accounts of the data.

These results suggest that (unsurprisingly), concrete rewards are the singular biggest contributor to subjective happiness. Across all the models we considered (despite their number of parameters), models that did not include a concrete reward term provided worse predictions. To avoid overwhelm and instead focus on evaluating the models that provide the best fits, we do not discuss models without concrete reward further.

### Models based on concrete rewards and token events (4 parameters)

This class of models focused on accounting for concrete rewards and 1 parameter that captured some aspect of token-related events. The best fitting model of this class incorporated both concrete rewards (CR) and token rewards (TR, model 4.1, mean r^2^=0.34). In the main text, we call this the “CRTR model”. An equivalent model based on CR and total token number provided a slightly worse account of the data (TT, model 4.2, mean r^2^=0.31).

When compared with 3 parameter models, these results suggest that both concrete and token rewards contribute to happiness fluctuations.

### Models based on concrete rewards and a decomposition of token-related events into 2 terms (5 parameters)

This class of models broke down token-related events into two terms, formulated in different ways. One best fitting model of this class parsed token events into tokens received when the certain reward (CrR) versus the gambling (GR) option was selected (as suggested by (2), model 5.1, mean r^2^=0.37). Similar fits were obtained when tokens were decomposed into expected value (EV2) and reward prediction error (RPE2), computed as described by (4), where EV2 was computed as the average of all token rewards up to the trial in question and RPE2 was computed as the difference between the tokens received and that value (model 5.3, mean mean r^2^=0.37). In comparison, a model that decomposed the total number of tokens into token reward (TR) and tokens at the start of each trial (TS) did not perform quite as well (model 5.2, mean r^2^=0.34).

These results suggest that happiness depends not just generically on the numbers of tokens won or lost on each trial. Rather, models that allow for happiness to be differentially weighted when (e.g.) the tokens won on each trial were won when the certain reward versus the gambling option was selected provide better fits.

### Models based on concrete rewards and a decomposition of token-related events into 3 terms (6 parameters)

This class of models focused on accounting for concrete rewards and token events parsed into 3 terms, including 1 term corresponding to tokens received when the certain reward (CrR) option was selected and 2 terms corresponding to tokens received on gambling trials. Events on gambling trials parsed two ways produced equivalent model fits. The first defined expected value (EV3) similar to model 5.3: as the average outcome up to each trial in question but limited in this case to gambling trials, following on ((4), model 6.1, mean r^2^=0.38). In the main text, we call this the “CRTE model”. The second defined expected value (EV1) as the average between the two options on gambling trials, following on ((2), model 6.2, mean r^2^=0.38). In both cases, reward prediction error (RPE3 and RPE1) was computed as the difference between the gambling trial outcome and EV.

#### General conclusions

Two 5 parameter model and two 6 parameter models provided nearly equivalent, best fits overall (model 5.1 and 5.3 mean r^2^ = 0.37; model 6.1 and 6.2, mean r^2^ = 0.38). Likewise, performance of the best fitting 4 parameter model was slightly but not markedly less (model 4.1, mean r^2^ = 0.34). Given the similarity of model fits, coupled with the similarities in the parameterization of these models, we hesitate to declare that there is a singular “best model” that accounts for subjective happiness reports in this data. Rather, a more parsimonious description is that a class of models in which happiness depends on a weighted combination of concrete rewards and token-related events is a decent description of the data overall, and breaking token-related events into event subtypes generally accounts for additional variance (with some nuances). For the clarity of presentation in the main body of the paper, we describe our approach to the neural data by focusing on a combination of A) model 4.1 (CRTR, where happiness depends on concrete and token reward, mean r^2^ = 0.34) due to the conceptual simplicity of that model and B) model 6.1 (CRTE, where happiness depended on concrete rewards and token reward broken into 3 terms) to illustrate the properties of the class of models with the best predictions. We also explore how behavioral and neural correlations compare across all models (Figure 4).

### Comparison of monkey choice models

**Supplemental Table 3.**
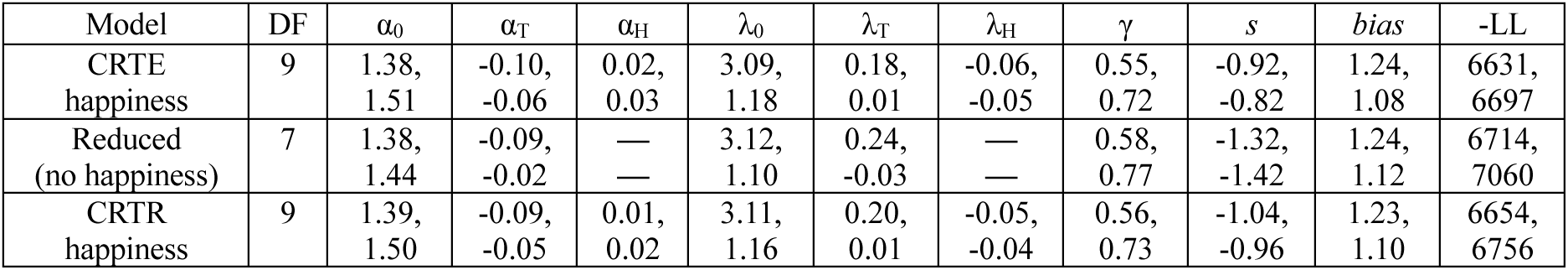
The table summarizes for each model the likelihood maximizing (best) parameters fit to data across sessions and its fitting performance for each monkey (within each cell, M1 top row; M2 bottom row), shown (as rows): when happiness was computed with the CRTE model, a reduced version of the model with no happiness terms, and when happiness was computed with the CRTR model. See Methods for a description of model parameters. See Results for cross-validated model comparison statistics. Final column: negative log likelihood (-LL) of models fit to all data. Comparing the reduced model with the CRTE model (lratiotest): M1, χ^2^(2)=164.5, P<0.001; M2, χ^2^(2)=726.1, P<0.001). Comparing the reduced model with the CRTR model (lratiotest): M1, χ^2^(2)=118.5, P<0.001; M2, χ^2^(2)=607.4, P<0.001).

| Model | DF | $\alpha_0$ | $\alpha_T$ | $\alpha_H$ | $\lambda_0$ | $\lambda_T$ | $\lambda_H$ | $\gamma$ | $s$ | *bias* | -LL |
| --- | --- | --- | --- | --- | --- | --- | --- | --- | --- | --- | --- |
| CRTE<br>happiness | 9 | 1.38,<br>1.51 | -0.10,<br>-0.06 | 0.02,<br>0.03 | 3.09,<br>1.18 | 0.18,<br>0.01 | -0.06,<br>-0.05 | 0.55,<br>0.72 | -0.92,<br>-0.82 | 1.24,<br>1.08 | 6631,<br>6697 |
| Reduced<br>(no happiness) | 7 | 1.38,<br>1.44 | -0.09,<br>-0.02 | —<br>— | 3.12,<br>1.10 | 0.24,<br>-0.03 | —<br>— | 0.58,<br>0.77 | -1.32,<br>-1.42 | 1.24,<br>1.12 | 6714,<br>7060 |
| CRTR<br>happiness | 9 | 1.39,<br>1.50 | -0.09,<br>-0.05 | 0.01,<br>0.02 | 3.11,<br>1.16 | 0.20,<br>0.01 | -0.05,<br>-0.04 | 0.56,<br>0.73 | -1.04,<br>-0.96 | 1.23,<br>1.10 | 6654,<br>6756 |

